# Place cell map genesis via competitive learning and conjunctive coding in the dentate gyrus

**DOI:** 10.1101/748640

**Authors:** Soyoun Kim, Dajung Jung, Sébastien Royer

**Affiliations:** Center for Functional Connectomics, Korea Institute of Science and Technology, Seoul, 02792, Republic of Korea; Center for Neuroscience Imaging Research, Institute for Basic Science (IBS), Suwon 16419, Republic of Korea; Department of Biological Sciences, Korea Advanced Institute of Science and Technology, Daejeon 34141, Republic of Korea; Division of Bio-Medical Science and Technology, KIST School, Korea University of Science and Technology, Seoul, 02792, Republic of Korea

**Author notes:** **Correspondence:** Sébastien Royer.

## Abstract

Place cells exhibit spatially selective firing fields and collectively map the continuum of positions in environments; how such network pattern develops with experience remains unclear. Here, we recorded putative granule (GC) and mossy (MC) cells from the dentate gyrus (DG) over 27 days as mice repetitively ran through a sequence of objects fixed onto a treadmill belt. We observed a progressive transformation of GC spatial representations, from a sparse encoding of object locations and periodic spatial intervals to increasingly more single, evenly dispersed place fields, while MCs showed little transformation and preferentially encoded object locations. A competitive learning model of the DG reproduced GC transformations via the progressive integration of landmark-vector cells and grid cell inputs and required MC-mediated feedforward inhibition to evenly distribute GC representations, suggesting that GCs progressively encode conjunctions of objects and spatial information via competitive learning, while MCs help homogenize GC spatial representations.

## Introduction

Principal cells in the hippocampus exhibit spatially selective firing fields called ‘place fields’ that collectively map the continuum of positions in environments (O’Keefe and Dostrovsky, 1971). How such activity patterns emerge in the DG, the first processing stage of the hippocampal trisynaptic loop, is a fundamental question. GCs account for nearly half of the neurons in the mammalian hippocampus (van Dijk et al., 2016); locally interact with MCs, a small population of excitatory neurons located in the hilus (10^4^ MCs versus 10^6^ GCs in rodents (Amaral et al., 1990; Amaral and Lavenex, 2007; Boss et al., 1985)); and are largely believed to perform pattern separation (Knierim and Neunuebel, 2016; Marr, 1971; McHugh et al., 2007; Rolls and Kesner 2006; Treves and Rolls, 1994) via the generation of sparse-orthogonal activity patterns (Danielson et al., 2017; GoodSmith et al, 2017; Hainmueller et al., 2018; Leutgeb et al., 2007; Senzai and Buzsáki 2017) and to assist memory formation in the CA3 via powerful GC-to-CA3 synapses (Amaral et al., 1990; Amaral and Lavenex, 2007; Henze et al., 2002; Rolls and Kesner, 2006; Treves and Rolls, 1994). Several mechanisms might be involved in GC place field generation.

First, object and spatial information might be integrated in the DG (Kesner et al., 2015; Rolls and Kesner, 2006). Inputs from both the lateral (LEC) and medial (MEC) divisions of the entorhinal cortex (EC) converge onto GCs (Amaral and Lavenex, 2007). Given that ‘landmark-vector’ cells (or ‘object-vector’ cells) in the MEC (Hoydal et al., 2019) and LEC (Deshmukh and Knierim, 2011) encode animal spatial relationships with objects and that ‘grid cells’ in the MEC exhibit periodic firing fields that convey spatial information related to path integration (Hafting et al., 2005; McNaughton et al., 2006), GCs might integrate object and spatial information via the integration of grid cell and landmark-vector cell inputs. Second, GCs are thought to operate as a competitive network in which strong feedback inhibition favors a ‘winner-take-all’ process and allows only a few GCs to be simultaneously active (Rolls et al., 2006; Si and Treves 2009). Interestingly, single place field representations develop naturally from the recursive updating of synaptic weights by Hebbian synaptic plasticity mechanisms in competitive network models, a process referred to as ‘competitive learning’ (Rolls et al., 2006; Si and Treves 2009). Third, MCs receive direct inputs from the CA3, semilunar GCs and possibly the EC (Larimer and Strowbridge, 2010; Scharfman, 2016) and are particularly well positioned to shape GC activity via both direct and indirect connections. In particular, MC-to-GC feedforward inhibition, the predominant contribution of MCs (Buzsáki and Czéh 1981; Scharfman 1995, 2016), might significantly influence competitive learning in the GC network.

To test these hypotheses, we recorded putative GCs and MCs as mice ran on a long treadmill enriched with visual-tactile cues (Royer et al., 2012), an apparatus facilitating the differentiation of spatial mechanisms (Fattahi et al., 2018; Geiller et al., 2017). We analyzed how spatial representations evolved as mice became familiarized with the belt layout over several days and, subsequently, how cells encoded other belt layouts. Finally, using computational modeling, we explored the network mechanisms and parameter conditions critical to reproduce the data. Our findings suggest that GC spatial representations develop over several days via competitive learning, encode conjunctions of grid cell and landmark-vector cell inputs, and evenly map the treadmill belt by means of an increase in MC-mediated feedforward inhibition in the cue locations.

## Results

### Identification of putative GCs and MCs

We recorded neuronal activity during treadmill running every day for 27 days using a 6-shank silicon probe (64 channels) implanted in the DG of the right brain hemisphere (Figure 1A and B; Figure S1). A total of 4003 cells were isolated (from 4 mice, 16 sessions per mouse) following standard criteria for unit detection and clustering (Harris et al., 2000; Kadir et al., 2014). In addition, to help identify putative GCs and MCs, we used a previous data set in which a subset of GCs and MCs expressed the Chronos opsin (via AAV/hSyn-Flex-Chronos-GFP injections in POMC-Cre and DRD2-Cre mice, respectively) and showed excitatory responses to light stimuli (18 POMC light-excited cells from 3 mice and 33 DRD2 light-excited cells from 2 mice) (Jung et al., 2019).

**Figure 1.**
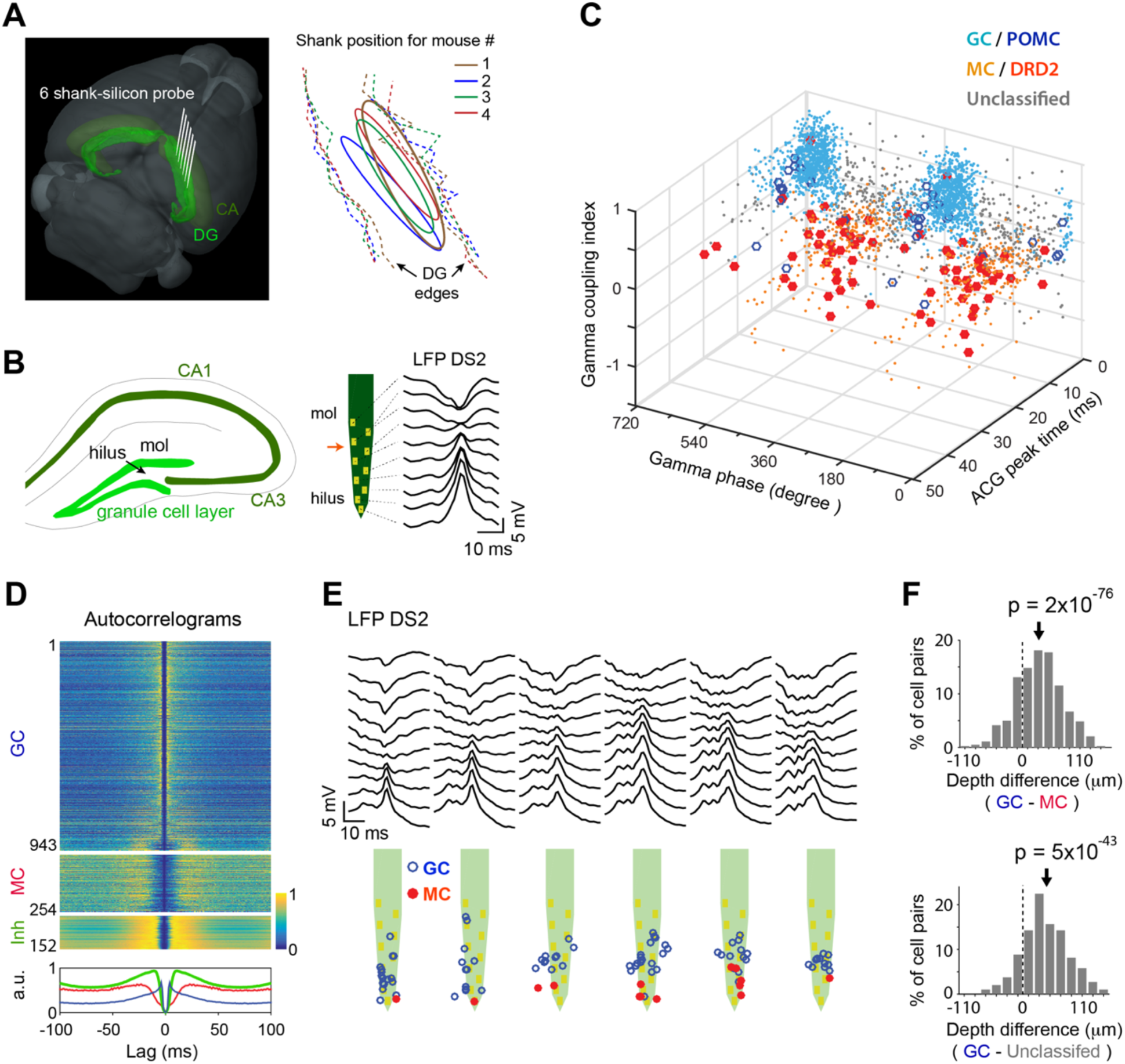
Recording of putative GCs and MCs. A. *Left*, 3D representation of the mouse brain (Allen Mouse Brain Institute; www.alleninstitute.org) showing recording electrode configuration in the dentate gyrus (light green). Dark green, cornus ammonis (CA). *Right*, the electrode positions (ellipsoid) relative to the lateral/medial edges of the granule cell layer (dashed lines) for all mice. B. *Left*, scheme showing the location of the hills and granule cell layer on a coronal section of the hippocampus. *Right*, layout of recording sites for a shank of the silicon probe and profile of local field potential dentate spike 2 (LFP DS2). Red arrow, position of DS2 reversal. C. 3D scatter plot for cells’ spike gamma phase, ACG refractory gap and gamma coupling index. Putative MCs (*orange dots*) and GCs (light *blue dots*) are identified by overlap with DRD2 (*red filled circle*) and POMC (*blue circle*) light-excited cells. D. Spike ACGs (*upper*, color-coded representation of individual cell; *lower*, population average) for putative GCs (*blue*), MCs (*red*) and inhibitory cells (*green*). E. Layout of LFP DS2 (*upper*) and putative GCs and MCs (*lower*) along the silicon probe shanks for one session. F. *Top*, depth differences between putative GCs and MCs recorded concurrently on the same shanks (*arrow*, the mean; paired t-test). *Bottom*, same analysis between GCs and the population of unclassified cells in panel C.

To allow the identification of putative GCs and MCs based on physiological criteria, we searched for differences in spike features between POMC and DRD2 light-excited cells. The features that best separated the two populations were cells’ spike autocorrelogram (ACG) and cells’ spike relationship with hilar local field potential gamma (30-80 Hz) oscillations (Jung et al., 2019) (Figure S2A-D): Spike ACGs were characteristic of short interval burst activity for POMC light-excited cells and showed large humps flanking a wide refractory gap in DRD2 light-excited cells, consistent with GC and MC ACGs observed during *in vivo* intracellular recordings (Henze and Buzsáki, 2007); POMC light-excited cells showed a preference to discharge near the troughs of gamma oscillations (Figure S2B) and were positively coupled with gamma power (Figure S2C), while DRD2 light-excited cells showed no clear gamma phase preference and were negatively coupled with gamma power. To quantify these differences, for individual cells, we implemented an ‘ACG refractory gap’ measure, defined as the duration for the ACG to reach 75% of its peak value (Figure S2A), and a ‘gamma coupling index’, defined as the difference in gamma power between window periods within (−10 to +10 ms) and outside (+40 to +100 ms) epochs of maximal firing activity (Figure S2D).

To identify putative GCs and MCs in the new data set, we measured the ACG refractory gap, gamma coupling index and mean gamma phase for each cell and examined the cell clustering and overlap with POMC/DRD2 light-excited cells and putative excitatory neurons (detected from short-latency peaks in cell-pair cross-correlograms (Barthó et al., 2004)). First, some cells were categorized as putative interneurons (n = 248) based on their high firing rate, short ACG refractory gap and lack of overlap with putative excitatory neurons (Figure S3A), and were excluded from the next analysis. Then, putative GCs (n = 2323) were characterized by short ACG refractory gaps, high gamma coupling indexes, a preference to discharge before the troughs of gamma oscillations, and overlap with POMC light-excited cells (Figure 1C and D; Figure S3B-E). In contrast, putative MCs (n = 408) were characterized by large ACG refractory gaps, low gamma coupling indexes, a preference to discharge before the peaks of gamma oscillations, and overlap with DRD2 light-excited cells. Consistent with anatomical figures, putative GCs were located closer to the reversal point of LFP type-2 dentate spikes (DS2) above the granule cell layer, while putative MCs were relatively deeper in the positive phase of DS2, i.e., in the hilus (Bragin et al., 1995) (Figure 1E); accordingly, on shanks where both putative GCs and MCs were detected, MCs were located on average 31.5±1.5 µm below GCs (t_824_ = 20.6, p = 2e^-76^, two-tailed paired t-test, n = 47 shanks, average over all GC-MC combinations; Figure 1F, top). Furthermore, a group of unclassified cells showing short ACG refractory gaps similar to those of GCs and similar gamma relationships to those of MCs (n = 168; Figure S3B) were found on average 43.8**±**2.7 µm below GCs (t_300_ = 16.2, p = 5e^-43^, two-tailed paired t-test, n = 13 shanks; Figure 1F, bottom), suggesting these cells might be pyramidal cells from CA3. Finally, a number of cells (n = 370) were not included in this classification because the number of spikes discharged was too low to reliably measure ACG refractory gaps and gamma phases (see Methods).

### Progressive transformation of GC firing fields over days

To investigate place cell activity during familiarization with the belt layout, mice were trained to run head-fixed for a water reward on an empty 150-cm-long belt for a week and then were introduced to a 201-cm-long belt displaying 3 pairs of visual-tactile landmarks (Figure 2A). The landmarks consisted of 5-cm-long arrays of small erect objects (shrink tubes, Velcro pieces, or glue spines) that lined both edges of the belt and provided visual-tactile stimulation to both sides of the mice. A water reward was delivered through a lick port on every trial (belt cycle) at the same belt position.

**Figure 2.**
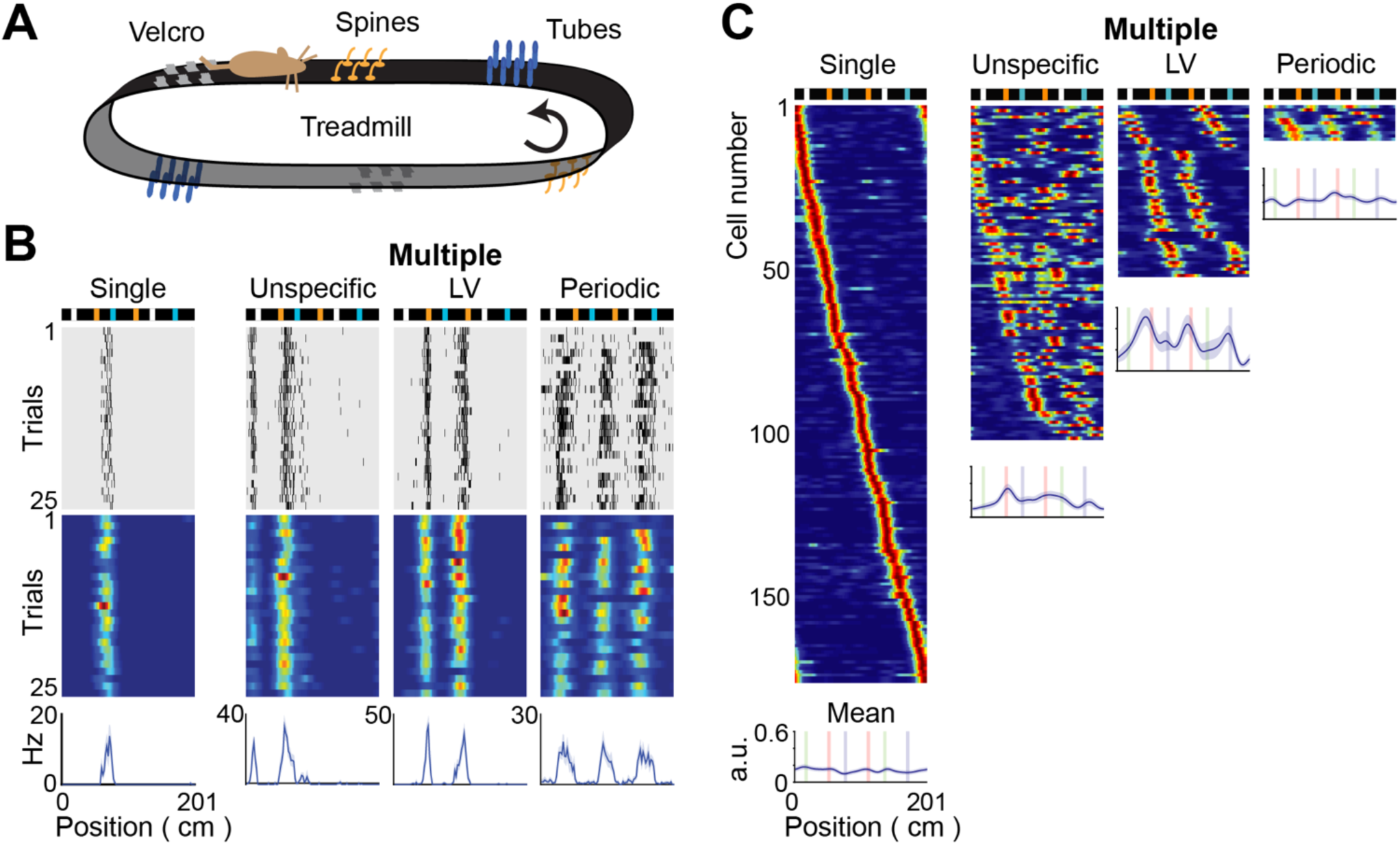
Types of place field activity among GCs. A. Scheme of the treadmill showing the 3 pairs of landmarks fixed on the belt. B. Individual cell examples for various types of GC representations: a single field cell; a landmark-vector (LV) cell (showing 2 fields matching a type of landmark); a periodic cell (with 3 periodic fields); and an unspecific cell (with more than 1 field that are neither LV nor periodic). *Top*, scheme of the belt; *middle*, spike raster and color-coded firing rate map; *bottom*, mean firing rate. C. *Color-coded*, firing rate maps of GCs from all sessions, sorted according to field positions and grouped by type of representation. *Line plots*, average of firing rate maps for each type of representation (*lines*, the mean; *shadows*, s.e.m).

We distinguished 4 types of place field activity among GCs (Figure 2B and C): 1) single place field cells; 2) landmark-vector (LV) cells, which, similar to LV cells in the EC and CA1 (Geiller et al., 2017; Hoydal et al., 2019), encoded spatial relationships with landmarks; 3) periodic cells, which exhibited 3 periodic firing fields with similar periodicity but various offsets, reminiscent of grid cells in the EC (Hafting et al., 2005) (interestingly, only a spatial period matching a third of the belt length was observed, in contrast to the multiple periods that grid cells encode, which might be because such a spatial period, equal to a submultiple of the belt length, enabled a spatially consistent recurrence of the firing fields across trials); and 4) ‘unspecific’ cells, which exhibited several firing fields but were neither LV cells nor periodic cells.

GC representations gradually transformed across days (Figure 3A-C). On day 1, a few GCs exhibited place fields (1.6±0.5% single and 5.8±2.2% multiple field cells (n = 4)) and were mostly LV cells and periodic cells (Figure 3B and C; among multiple field cells, 43.3±19.0% LV, 44.4±22.2% periodic and 12.2±6.2% unspecific cells). Then, the fraction of GCs with place fields progressively increased via an increase in unspecific and single field GCs, reaching a plateau after 5 and 10 days, respectively, while the fraction of LV cells and periodic cells decreased (Figure 3B and C; for days 13-20, 20.2±3.3% single and 11.8±1.6 multiple field cells (n = 12); among multiple field cells, 21.0±5.3% LV, 1.7±1.7% periodic and 77.4±5.6% unspecific cells). Moreover, the peak firing rates of the two fields of LV cells became increasingly uneven (Figure 3D; n = 53, r = -0.5, p = 1e^-4^, Pearson’s correlation; peak rate ratio on day 1 versus days 13-20, 81.8±5.8% (n = 6) versus 36.8±5.3% (n = 10), t_14_ = 5.5, p = 8e^-5^, two-tailed unpaired t-test).

**Figure 3.**
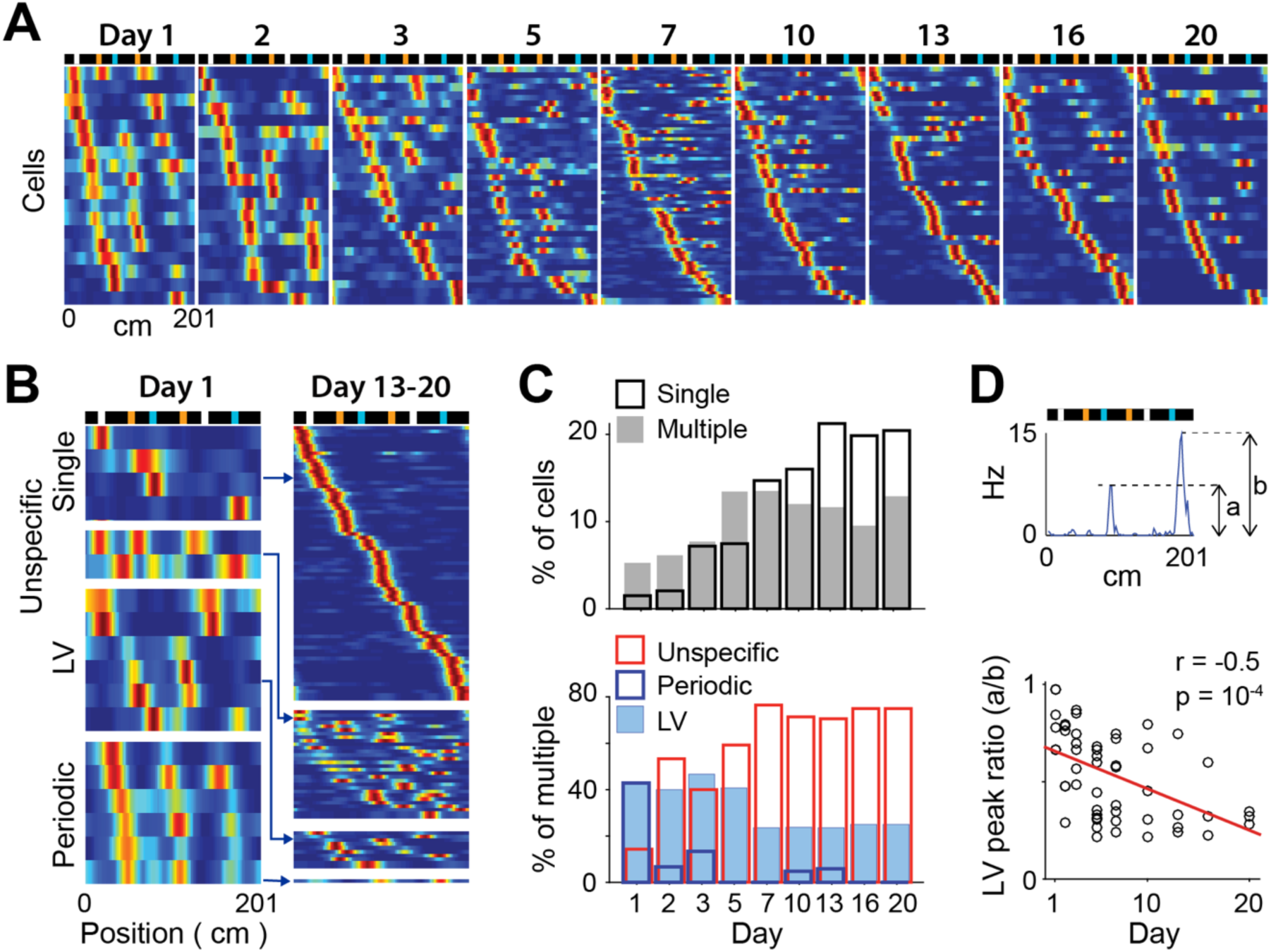
Progressive transformation of GC firing fields across days. A. Color-coded firing rate maps for GCs showing firing fields across days. The rows of the matrices correspond to individual GCs and are sorted according to firing field positions. *Top*, scheme of the belt. B. Color-coded firing rate maps of GCs on day 1 (left) and on days 13, 16 and 20 combined (right), for each type of GC firing field. C. *Upper*, proportion of GCs with a single field (black) and multiple fields (gray) across days. *Lower*, proportion of LV, periodic and unspecific GCs, among multiple field GCs, across days. D. *Upper*, definition of LV peak ratio as the ratio between LV fields’ peak firing rates (smaller peak over larger peak). *Lower*, distribution of LV peak ratios across days. Each dot is the LV peak ratio of one LV cell. *Red line*, linear fit. n = 53, r = -0.5, p = 1e^-4^, Pearson’s correlation.

### Emergence and extinction of firing fields within sessions

The increase of single field representations and the reduced proportion of LV and periodic cells implies that new place fields emerged and that existing place fields became extinct. Both place field emergence and extinction events could be observed within sessions (Methods, Figure 4A) and produced preferentially incremental changes in the number of firing fields in each cell (Figure 4B, F_1,48_ = 4.5, p = 0.04, two-way ANOVA). Place field emergences were characterized by a steep rise of the in-field rate to a plateau value (Figure 4C) and occurred preferentially at the beginning of the sessions (Figure 4D; percent of events before versus after trial 30, 75.6±10.2% versus 24.4±10.2%, t_15_ = 2.5, p = 0.02, two-tailed paired t-test), while place field extinctions were preceded by gradual decreases in the in-field firing rate (Figure 4C) and mostly occurred late in the sessions (Figure 4D; percent of events before versus after trial 30, 20.8±8.9% versus 79.2±8.9%, t_15_ = -3.3, p = 5e^-3^, two-tailed paired t-test).

**Figure 4.**
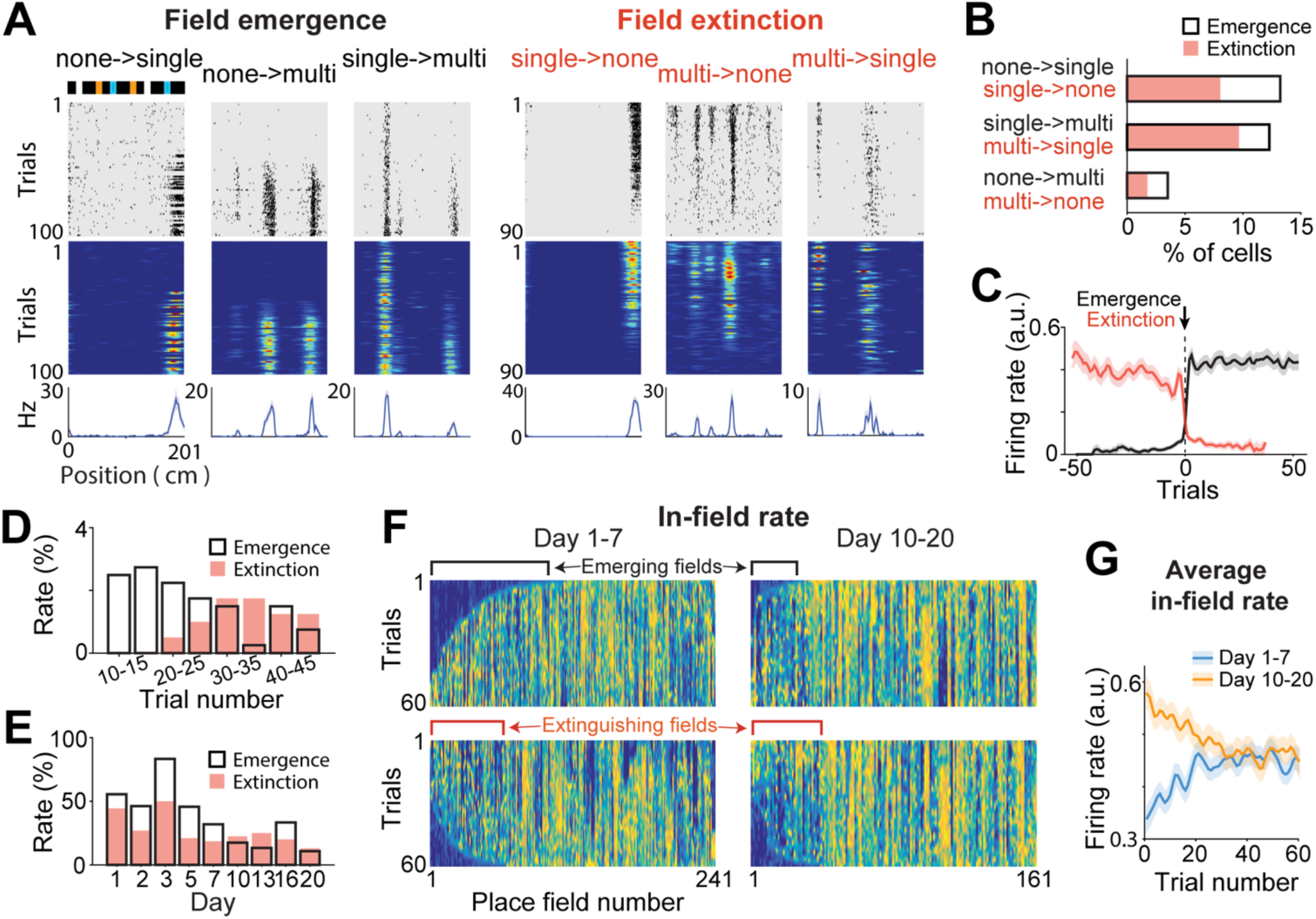
Emergence and extinction of firing fields within sessions. A. Individual cell examples for field emergences and field extinctions within a session for GCs converting between no field, single field and multiple field conditions. *Top*, scheme of the belt; *middle*, spike raster plot and color-coded firing rate map; *bottom*, mean firing rate. B. Proportion of GC conversion types for field emergences (black) and field extinctions (salmon). Proportions are relative to the number of GCs with place fields. C. Average in-field firing rate (lines, the mean; shadow, s.e.m) for field emergences (black) and field extinctions (salmon), after aligning to emergence (or extinction) trials of in-field firing rate vectors. Note that field emergence is relatively instantaneous, as the in-field firing rate reaches an immediate plateau, while field extinction is preceded by a gradual decrease in the in-field firing rate. D. Proportion of field emergence (black) and field extinction (salmon) events as a function of session trials. E. Proportion of field emergence (black) and field extinction (salmon) events as a function of days. F. Color-coded representation of in-field firing rate vectors for all place fields on days 1-7 (left) and days 10-20 (right). In-field firing rate vectors are sorted to emphasize field emergences (*upper*, sorted by trial number for which cumulative sums > 20% of vector integrals) and field extinctions (*lower*, using the same sorting method in the reversed direction). G. Average in-field firing rate (lines, the mean; shadow, s.e.m) over trials for days 1-7 (blue) and days 10-20 (yellow).

Importantly, emergence and extinction rates changed across days, which was consistent with the gradual transformation of GC representations. While both emergence and extinction rates decreased across days (Figure 4E; F_8,42_ = 3.7, p = 2e^-3^, two-way ANOVA, emergence rate across days 1-7 versus days 10-20, 43.1±6.6% versus 19.9±5.1%, t_24_ = 2.8, p = 0.01, two-tailed unpaired t-test; extinction rate, 34.6±5.6% versus 21.5±3.5%, t_24_ = 2.0, p = 0.06, two-tailed unpaired t-test), the emergence rate was initially higher than the extinction rate and reached an equivalent level after 7 days (emergence versus extinction, days 1-7, t_4_ = 5.0, p = 7e^-3^; days 10-20, t_3_ = -0.26, p = 0.81, two-tailed paired t-test), matching the increase in and stabilization of place cells observed in Figure 3C. This effect was also observable in the matrix concatenation of in-field firing rates for all GC place fields, sorted by time of field emergence or extinction (Figure 4F) and was also revealed by distinct profiles of average in-field rate for days 1-7 and 10-20 (Figure 4G).

### Changing the belt

The gradual transformation of GC representations might be associated with the development of an engram specific to the particular features of the belt. To test the dependence of place cell activity on belt features, after day 21, we recorded the same neurons across 3 consecutive sessions using 3 distinct belt layouts: the ‘original’ belt layout; a ‘reordered’ belt, presenting the same landmarks as the original belt but in a rearranged order; and a ‘novel’ belt, which was a different length (211 cm) and presented a new set of landmarks (Figure 5A).

**Figure 5.**
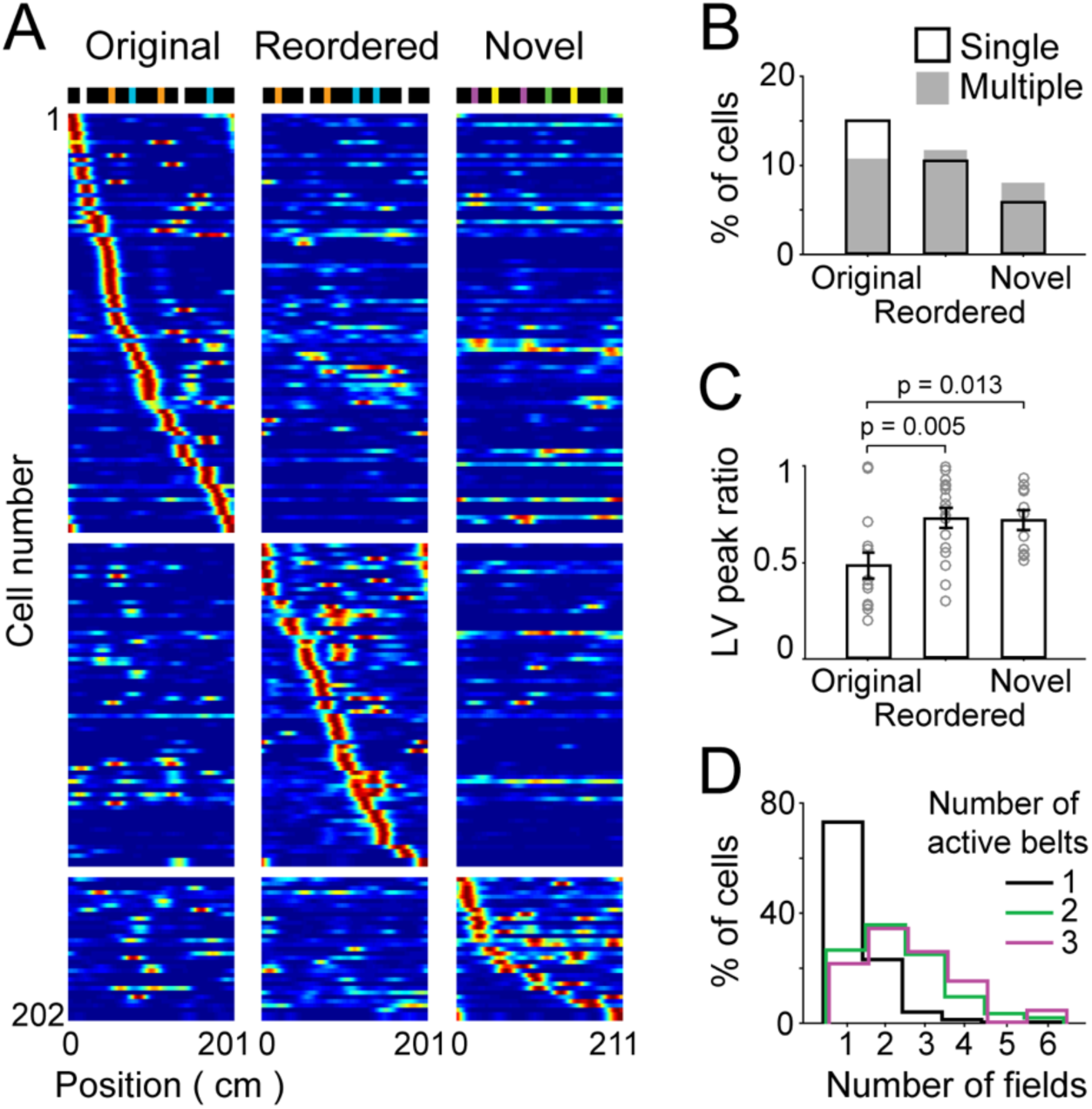
GC encoding of other belt layouts. A. Firing rate maps of GCs for the ‘original’ (left), ‘reordered’ (center) and ‘novel’ (right) belts. *Top*, scheme of the belt layouts. *Color-coded*, each row represents the firing rate maps of a GC for the 3 belts, normalized by the peak firing rate of the cell across all belts. Only GCs exhibiting place fields in at least one of the belts are displayed. GCs are sorted according to the type of belt and the belt position of the largest firing fields. B. Fraction of GCs with single (black) and multiple (gray) place fields for each belt.. C. LV peak ratio (*circle*, individual cell; *bar*, the mean; error bar, s.e.m) of LV cells for the 3 belts. Note that the magnitude of the two firing fields is more similar in the reordered and novel belts than in the original belt (F_2,40_ = 6.2, p = 5e^-3^, one-way ANOVA; t_30_ = -3.0, p = 5e^-3^, t_24_ = -2.7, p = 0.01 respectively, ad hoc two-tailed unpaired t-test). D. For the groups of GCs that are active in 1 (black), 2 (green) and 3 (purple) belts, the distribution of the number of fields per cell (in the belt with the largest number of fields). Note that cells exhibiting multiple place fields tend to represent several belts, while cells exhibiting a single place field tend to represent only one belt, t_215_ = 7.2, p = 1e^-11^, two-tailed unpaired t-test).

Consistent with the idea that an engram specific to the layout of the original belt was created, GCs exhibiting single place fields were less frequent in sessions using the reordered and novel belts than in sessions using the original belt (Figure 5B; original belt,15.0±1.3; reordered belt, 11.5±1.7%; novel belt, 7.1±1.5%; F_2,72_ = 10.47, p = 1e^-4^, three-way ANOVA; original vs reordered, t_27_ = 1.7, p = 0.10; original vs novel, t_27_ = 5.2, p = 2e^-5^, ad hoc two-tailed paired t-test; average across days 21-27); for LV cells, the magnitude of the two firing fields was more similar in sessions using the reordered and novel belts than in sessions using the original belt (Figure 5C; peak rate ratio, original belt, 0.49±0.06 (n = 15); reordered belt, 0.73±0.05 (n = 17); novel belt, 0.72±0.05 (n = 11); F_2,40_ = 6.2, p = 5e^-3^, one-way ANOVA; original vs reordered, t_30_ = -3.0, p = 5e^-3^, original vs novel, t_24_ = -2.7, p = 0.01, ad hoc two-tailed unpaired t-test; average across days 21-27). However, the fraction of single field cells was higher in sessions using the reordered belt than in those using the new belt (t_27_ = 2.5, p = 0.02, ad hoc two-tailed paired t-test), suggesting that the engram may have helped place field generation for other belts according to the degree of belt similarity.

Furthermore, a relationship was apparent between the number of place fields and the number of belts represented by each cell (Figure 5D). Cells exhibiting multiple place fields tended to show place fields for several belts, while cells exhibiting single place fields tended to be active in only one belt (average number of belts represented by multiple field cells, 2.03±0.07 (n = 110) and by single field cells, 1.35±0.06 (n = 107); t_215_ = 7.2, p = 1e^-11^, two-tailed unpaired t-test). Given that young adult-born GCs are more excitable, display more place fields and differentiate less the contexts (Danielson et al., 2016; Gonçalves et al., 2016; Marín-Burgin et al., 2012; Schmidt-Hieber et al., 2004), these findings are consistent with the idea that single and multiple field cells correspond to mature and young adult-born GCs, respectively.

### MCs show distinct representations

We distinguished 3 types of firing activity patterns among MCs: 1) cells with a single place field, 2) cells with multiple place fields, and 3) cells with relatively high firing rates (i.e., a mean firing rate > 3 Hz) but low spatial modulation (Figure 6A, see Methods). In contrast to GCs, the fraction of MCs showing firing activity on the belt was initially high (70.6±10.7% of MCs compared to 7.3±2.7% of GCs on day 1; t_3_ = 6.4, p = 8e^-3^, two-tailed paired t-test) and did not change significantly across days (Figure 6B; days 1-7 versus days 10-20, 67.2±3.3% (n = 20) versus 61.4±6.4% (n = 16); t_34_ = 0.9, p = 0.40, two-tailed unpaired t-test); and for all sessions, spatially modulated MCs showed mostly multiple firing fields (Figure 6B-D), even though the number of fields per cell decreased across days (Figure 6C; days 1-7 versus days 10-20, 4.44±0.24 versus 3.51±0.25 fields per cell; n = 52, 39 respectively, t_89_ = 2.8, p = 7e^-3^, two-tailed unpaired t-test).

**Figure 6.**
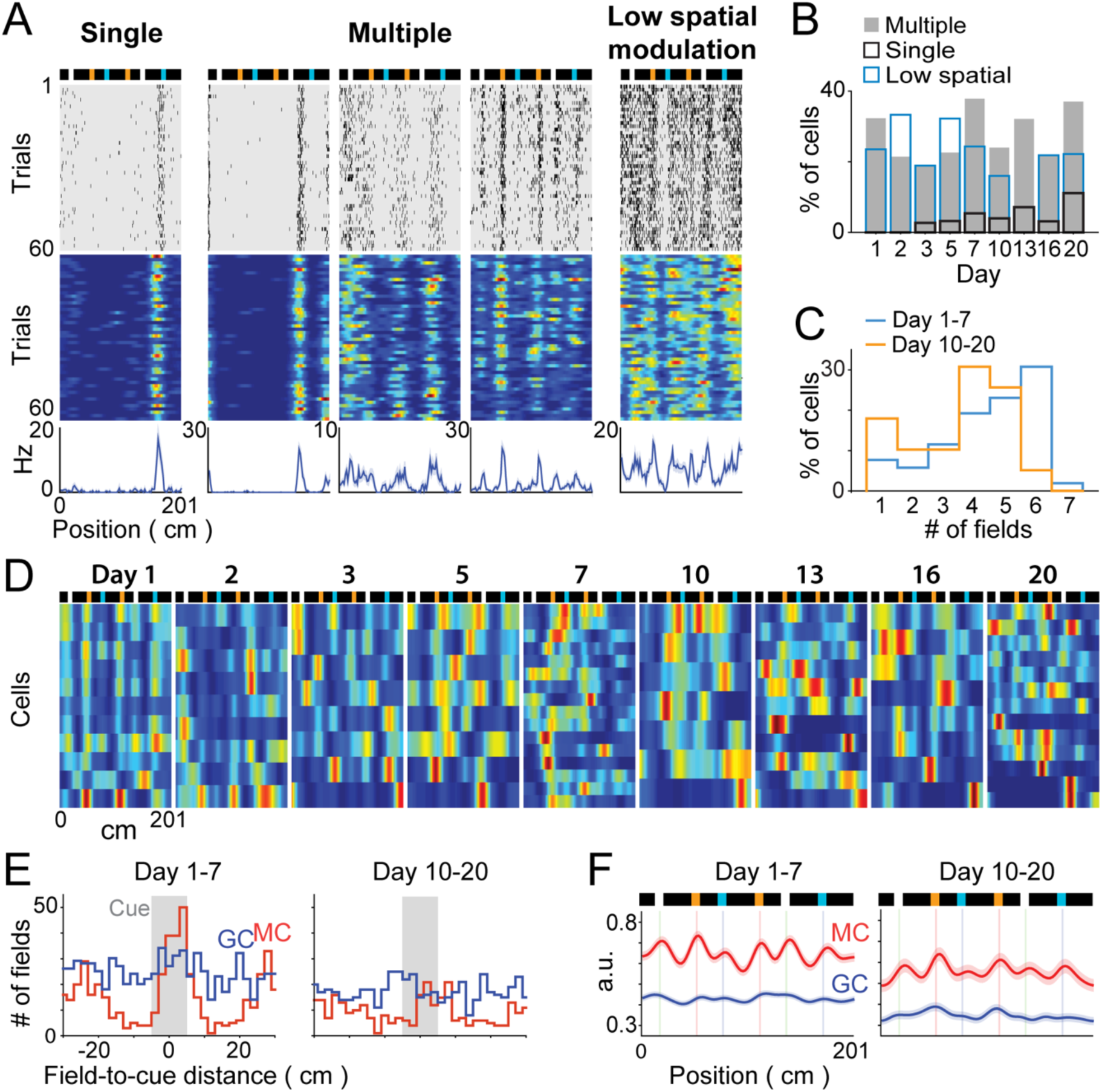
Distinct MC representations. A. Individual cell examples for various types of MC representations. From left to right, a single field cell, three multiple field cells, and a cell with low spatial modulation. *Top*, scheme of the belt; *middle*, spike raster plot and color-coded firing rate map; *bottom*, the mean firing rate. B. Proportion of MCs with single fields (black), multiple fields (grey) and low spatial modulation (blue), across days. C. Distribution of the number of fields per cell for days 1-7 and days 10-20 among MCs exhibiting place fields. The number of place fields per cell was reduced across days (t_89_ = 2.8, p = 7e^-3^, two-tailed unpaired t-test). D. Color-coded rate maps for MCs exhibiting firing fields across days. The rows of the matrices correspond to individual MCs and are sorted according to firing field positions. E. Distribution of field-to-landmark distances for GC (blue) and MC (red) place fields for days 1-7 (left) and days 10-20 (right). F. Average of GC (blue) and MC (red) rate maps (line, the mean; shadow, s.e.m) for days 1-7 (left) and days 10-20 (right). Note the increase in MC (but not GC) activity in landmark positions.

Unexpectedly, MC firing fields were strongly modulated by the landmarks, as they were repeated at multiple landmark positions with little spatial offset from the landmarks (Figure 6D). Accordingly, both the distribution of field-to-landmark distances and the averaged cell firing rate maps showed clear peaks in landmark positions for MCs, but not for GCs, an effect that was reduced but still prominent during later sessions (Figure 6E and F; fraction of fields with peak < 5 cm from landmarks, days 1-7, MCs, 70.2±5.0% (n = 19), GCs, 50.0±5.4% (n = 19), t_36_ = 2.8, p = 9e^-3^; days 10-20, MCs, 49.6±5.8% (n = 12), GCs, 43.4±4.3% (n = 16), t_26_ = 0.9, p = 0.39; days 1-7 versus days 10-20, MCs, t_29_ = 2.6, p = 0.01, GCs, t_33_ = 0.9, p = 0.40, two-tailed unpaired t-test). Hence, MC spatial activity was likely not generated by GC inputs (consistent with Senzai and Buzsáki, 2017) but rather by other afferents such as CA3, semilunar granule cells (Larimer and Strowbridge, 2010) or direct inputs from the EC (Scharfman 1995, 2016).

### Modeling the increase in GC single place fields

The DG has been modeled as a competitive network in which discrete place field representations are generated via competitive learning (Rolls et al., 2006; Si and Treves, 2009). To test whether competitive learning could produce the gradual increase in GC single fields and the decrease in LV and periodic cells, we implemented a model of DG in which 3000 GCs received excitatory inputs from 300 EC grid cells (with spatial periodicity similar to that of the periodic cells) and 300 EC LV cells and, in addition, were subjected to feedback inhibition (Figure 7A (*gray and black)*, Figure S4A Methods). For a given belt position, the excitation (E) received by a GC was the weighted sum of EC cell activity, the feedback inhibition (I) was proportional to the sum of GC activity, and the activity of a GC was equal to E-I (if E>I) or 0 (if E<I). EC-to-GC synaptic weights were initially allocated randomly according to a gamma distribution and then modified on each iteration by two operations: Hebbian synaptic potentiation proportional to the degree of GC-EC coactivation and normalization of the total synaptic weight for each GC^38-40^ (Bliss and Collingridge, 1993; Royer an Paré 2003; Turrigiano et al., 1998) (Figure 7B (*black*), Figure S4B).

**Figure 7.**
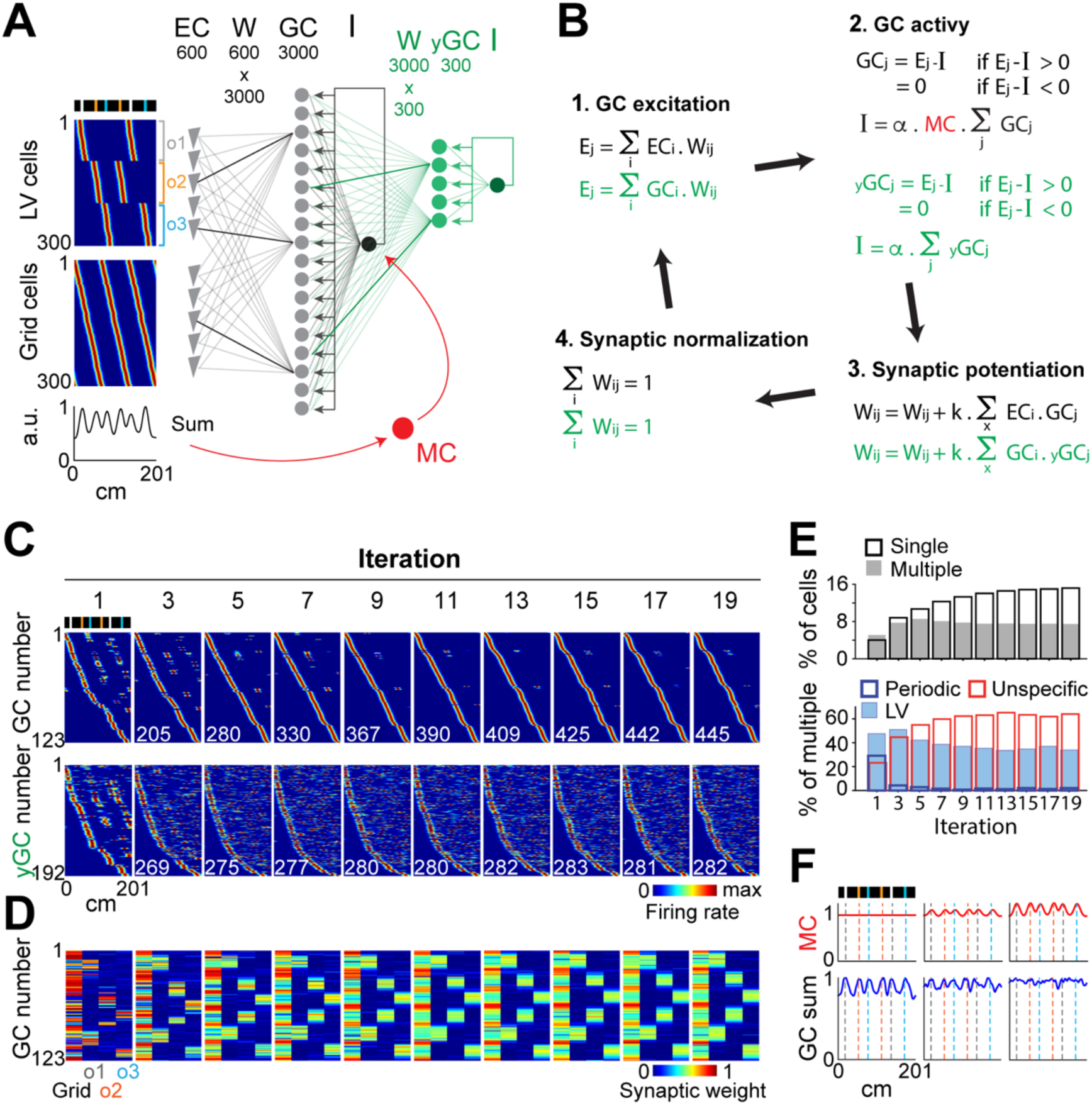
Competitive learning model. A. Model architecture. The simplest version of the model is shown in gray and black. 3000 GCs receive excitatory inputs from 300 LV cells and 300 grid cells from the EC, and are subjected to feedback inhibition. The EC-to-GC synaptic weight matrix is referred to as Wij. The threshold used as feedback inhibition is referred to as I. *Color coded*, rate maps of EC LV cells (upper) and grid cells (lower). LV cells encode various distances to landmarks, while grid cells have various spatial phases and the same periodicity as periodic cells. The first modification of the model is shown in green. Three hundred ‘immature’ GCs (yGCs) that receive inputs from the 3000 ‘mature’ GCs and that are subjected to a low level of feedback inhibition are added to the previous model. The second modification of the model is shown in red. MC feedforward inhibition is added to the previous model. The spatial modulation of MC activity is assumed to be proportional to the dynamic range of EC average activity. B. All operations were executed during one model iteration. 1) The excitation Ej received by the GCj in a given position. 2) The levels of GCj activation and feedback inhibition I in that position. The value of I is estimated numerically by finding the value among a range of I values that best satisfies the two relations. 3) The potentiation of synaptic weights, proportional to the level of cells’ co-firing throughout the belt. 4) The normalization of synaptic weights. C. Transformation of spatial representations across iterations. Color-coded representation of rate maps for active mature GCs (upper) and active immature GCs (lower), across iterations. Active cells have a mean activity > 0 and are sorted according to firing field positions. Note the increase in the number of active GCs (lower left numbers) and the transformation of representations from multiple to single place fields. Note also that immature GC representations initially resemble mature GC representations and then become multiple unspecific fields. D. For each active GC, the color-coded representation of the total synaptic weight contributed by grid cells and by each population of LV cells (encoding a specific landmark) is shown across iterations, using the same GC ordering as in c. Note that synaptic inputs initially originate from one source and are progressively redistributed equally among grid cells and LV cells. E. *Upper*, proportion of GCs (irrespective of type) with a single field (black) and multiple fields (grey), across iterations. *Lower*, proportion of LV, periodic and unspecific GCs, among multiple field GCs, across iterations. Note the similarity with experimental trends (Figure 3C). F. Effect of the distinct magnitude of MC modulation on the GC mean activity.

We found that two parameters critical to reproduce the gradual increase in GC single place field representations were the initial repartition of EC-to-GC synaptic weights and the strength of the feedback inhibition. For high levels of feedback inhibition and sparsity of initial inputs, the model could reproduce the initial preponderance of periodic and LV place fields and the asymptotic increase in single place field representations (Figure 7C, top; Figure S4C; Figure S5). When the sparsity of the initial inputs was set too low (i.e., each GC receives substantial excitation from several EC cells), an asymptotic decrease in single place field representations was observed over time; in contrast, when the sparsity of the initial inputs was set too high (i.e., each GC is mostly excited by one EC neuron), the fraction of cells that developed single place fields was too high. When the feedback inhibition was set too low, the fraction of cells with place fields (single, periodic or LV) was too high (Figure S5).

Importantly, the transformation of GC representations was accompanied by a specific reconfiguration of EC-to-GC synaptic weights. While most excitatory input to a GC was initially contributed by one EC neuron as a result of the initial sparse input setting, excitatory input to a GC was progressively shared by multiple EC neurons, and each GC progressively received, in comparable amount, inputs from grid cells and LV cells (Figure 7D; Figure S4C). Hence, this simple competitive learning model could reproduce the transformation of GC representations, from periodic and LV place fields to single place fields, through the progressive integration of grid cell and LV cell inputs.

### Modeling the increase in multiple unspecific GC place fields

The model did not, however, replicate the progressive emergence of multiple ‘unspecific’ place fields. Reducing feedback inhibition increased the number of multiple field cells, but these cells were mostly periodic or LV cells, and the overall fraction of cells generating place fields was excessive (Figure S5). Therefore, we reasoned that unspecific place fields should be generated by a small subset of GCs under low inhibition that receive inputs from cells other than EC grid cells and LV cells and that initially exhibit periodic and LV firing fields similar to other GCs. The small population of young adult-born GCs could match such criteria because they are generally more excitable and active than mature GCs, and they receive their first excitatory synapses from non-EC inputs, including mature GCs (Gonçalves et al., 2016; Schmidt-Hieber et al., 2004; Vivar et al., 2012). To test this hypothesis, we included a population of 300 young adult-born GCs under weak feedback inhibition that received excitatory inputs from mature GCs (Figure 7A (*green*), Figure S6) and that were subjected to the same synaptic plasticity mechanisms (Figure 7B). As anticipated, young adult-born GCs generated initially periodic and LV firing fields from the sparse inputs of mature GCs and then, over time, developed multiple unspecific place fields reflecting combinations of the single place fields of mature GCs (Figure 7C, bottom). As a result, the model could largely reproduce the experimental results for the ratio and temporal evolution of different cell representations (Figure 7E).

We next examined the predicted cell activity for the reordered and new belts. To simulate the reordered belt, the firing fields of EC LV cells were simply moved to the new location of the landmarks, whereas to simulate the new belt, EC LV and grid cells were randomly assigned to new object pairs and spatial grid phases, respectively (Figure S6E). Similar to the experimental data (Figure 5B and D), the number of single field representations was decreased for the reordered belt compared to the original belt and was further decreased for the new belt (Figure S6F), and the number of place fields per cell was correlated with the number of belts represented (Figure S6G). Hence, this version of the model could reproduce both the transformation of GC representations across days for the original belt and the subsequent encoding of the reordered and new belt layouts.

### Modeling the contribution of MCs to GC representations

Given that MC average activity was increased in landmark locations and that a predominant contribution of MCs is believed to be feedforward inhibition (Buzsáki and Czéh 1981; Scharfman 1995, 2016), we tested how an MC-mediated increase of inhibition in landmark locations affects GC representations (we considered only the network of mature GCs for this analysis). Since MCs presumably receive EC information both directly and indirectly (Scharfman 2016), we assumed that average MC activity reflects average EC activity and modulates the threshold parameter I of the model (Figure 7A and B (*red)*, Figure S7).

In contrast to the experimental findings, average GC activity progressively increased in landmark positions when the inhibition was not modulated by MCs (Figure 7F, Figure S7C). However, this effect decreased as MC modulation of inhibition was strengthened, such that GCs could evenly represent all belt locations (Figure 7F, Figure S7D-F). Hence, increasing inhibition in locations associated with larger excitation is necessary to achieve uniform mapping of the space via competitive learning, and a role of MC feedforward inhibition might be to enable the uniform mapping of the space by GCs.

## Discussion

Using a combination of silicon probe recording, treadmill running and modeling approaches, we were able to identify putative GCs and MCs, monitor the development of spatial representations for a particular layout of landmarks, observe subsequent encodings of other layouts, and interpret the results in terms of learning mechanisms, encoded information and cell-specific functions.

Our findings suggest that competitive learning underlies the increase in place field representations in DG. The model reproduced the transformation and the asymptotic increase in GC representations across days, as well as the subsequent encoding of other belt layouts. How place fields emerge through competitive learning is not straightforward, as cells that are silent paradoxically develop place fields as a result of Hebbian synaptic plasticity, which requires postsynaptic firing. In the model, the initial activation of silent GCs could only be elicited through disinhibition, which had to result from a local decrease in the mean GC activity following synaptic normalization. Hence, disinhibition induced by synaptic normalization might be the underlying mechanism for GC place field emergence during competitive learning. Interestingly, the development of GC spatial representations was relatively slow, plateauing after a week. Slow learning rates are theoretically critical for minimizing the alteration of prior memories by new learning (the stability-plasticity dilemma (Carpenter and Grossberg,1987)) and could reflect the necessity of sleep-dependent mechanisms. Periods of rest between treadmill running sessions were likely essential to regenerate the network potency for plasticity, as the place field emergence rate progressively decreased across trials within each session (Figure 4D). Synaptic normalization, especially, might require sleep and sharp-wave-ripple oscillations to develop (Norimoto et al., 2018; Vyazovskiy et al., 2008), considering the relatively slow rate of synaptic downscaling observed in vitro (Royer an Paré, 2003).

The model largely implied an integration of LV cell and grid cell inputs, such that GC activity was contingent upon the particular alignment of LV cells and grid cells on the belt. Such integration might underly the elaboration of memory engrams for spatial context (Hainmueller et al., 2018; Liu et al., 2012) comprising object location information within EC-to-GC synapses. Through such engrams, GCs would produce an output very specific to the spatial context and would remap when the object layout is changed (Jung et al., 2019), consistent with the importance of an intact DG to discriminate contexts and detect changes in object layouts (Hunsaker et al., 2008; Kesner et al., 2015; Rolls and Kesner, 2006). DG function might also be considered in terms of pattern separation, with the combination of context engram, sparse GC activity and inter-GC competition contributing to strong re-mapping responses by GCs (Jung et al., 2019). In this respect, our findings imply that pattern separation should improve with experience as context engrams progressively develop, whereas pattern separation in a new context should be enhanced by prior experience in other similar contexts (given the generalization of learning across similar belt layouts).

‘Unspecific’ GCs likely corresponded to immature GCs, as they exhibited multiple firing fields with low specificity to the context (Danielson et al., 2016), which is consistent with the high excitability of young adult-born GCs (Gonçalves et al., 2016; Marín-Burgin et al., 2012; Schmidt-Hieber et al., 2004). In this respect, the fact that unspecific GC firing fields could not be reproduced by the model with grid cell and LV cell inputs is consistent with reports that young adult-born GCs initially receive inputs mostly from MCs, the CA3 and mature GCs (Gonçalves et al., 2016; Vivar et al., 2012). Incorporating a population of young adult-born GCs that received mature GC inputs in the model was sufficient to reproduce unspecific GC firing fields, as the critical requirements were low feedback inhibition and excitation provided by single field cells instead of grid cells and LV cells. While the benefit of such a network configuration is unclear, it is possible that young adult-born GCs might implement in parallel pattern separation (via the mature GCs) and other operations requiring high cell excitability, such as the temporal binding of episodic information (Aimone et al., 2006). Furthermore, increasing inhibition in landmark locations was critical to reproduce the uniform mapping of the space by GCs, and MCs are well suited to support such an operation, as they showed firing rate increases in landmark locations and generate feed-forward inhibition in GCs (Buzsáki and Czéh 1981; Scharfman 1995, 2016). Hence, a function of MCs might be to sense variation in EC input through a parallel (non-GC) pathway, perhaps via CA3, semilunar GCs and/or direct EC afferents (Larimer and Strowbridge 2010; Scharfman 1995, 2016), and proportionally increase inhibition to ensure a uniform mapping of the space by GCs.

Our findings suggest that through competitive learning mechanisms and the integration of LV and grid cell inputs, the DG is particularly apt at generating firing fields that are spatially tuned, specific to contexts and stable in time (Hainmueller et al., 2018); however, this ability is at the cost of slow learning rates. While cells in other hippocampal regions might be more fit to encode dynamic features (Hainmueller et al., 2018) related to episodic memory, they might depend on DG output for both the spatial tuning and context specificity of firing fields (Lee et al., 2012) (consistent with the parallel decrease in GC number (Amaral and Lavenex, 2007) and CA3 spatial tuning (Royer et al., 2010) along the septo-temporal axis). However, the possibility remain that the DG might also form non-spatial representations in other circumstances, and whether competitive learning mechanisms operate on the whole DG network or on the level of DG subnetworks associated with distinct information, time scales or cell morphology (Diamantaki et al., 2016) needs to be examined in the future.

## Methods

### Animals

All experiments were conducted in accordance with institutional regulations (Institutional Animal Care and Use Committee of the Korea Institute of Science and Technology) and conformed to the Guide for the Care and Use of Laboratory Animals (NRC 2011). Four male C57BL6 mice between 6 and 7 weeks old were used. The mice were housed 2 per cage in a vivarium under a 12 hour light/dark cycle.

### Virus injection and preparation for head fixation

During a first surgery, under isoflurane anesthesia (supplemented by subcutaneous injections of buprenorphine 0.1 mg/kg, followed by daily subcutaneous injection of ketaprofen 5 mg/kg for 2 days), two small watch screws were driven into the bone above the cerebellum to serve as reference and ground electrodes for the recordings. A 3D printed plastic head-plate with a window opening in the center was cemented to the skull with dental acrylic. The head-plate was designed to be conveniently fixed/unfixed to a holding plate using 2 screws.

### Treadmill apparatus and behavioral paradigm

After a postsurgery recovery period of 7 days, the mice were water restricted to 1 ml of water per day and were trained for 2 weeks (one 1-hour session per day) to run on the treadmill with their heads fixed. The treadmill was not motorized but consisted of a light velvet belt laying on two 3D printed wheels, which mice moved themselves at will. Water rewards were delivered on every trial at the same position on the belt via a lick port. After behavioral learning reached an asymptote, the animals completed 100 to 150 trials in the first 45 min of each session. The quantity of water consumed on the treadmill was measured after each session, and additional water was provided such that the mice drank a total amount of 1 ml of water per day.

For recording sessions, landmarks were fixed and interspersed on a 211-cm long belt. The landmarks consisted of arrays of vertical poles and textures fixed on the edges of the belt, which provided visual-tactile stimulation to both sides of the mice.

### Chronic implantation of the electrode

Procedures for chronic recordings have been described previously (Chung et al. 2017; Geiller et al., 2017; Jung et al., 2019). Briefly, a craniotomy was performed under isoflurane anesthesia. A silicon probe was coated with red-fluorescent dye (DiI, Life technologies), mounted on a custom microdrive, and lowered into the DG, which was detected by the emergence of unit activity following an ∼500-µm silent zone below the CA1 pyramidal layer. The microdrive was then cemented to the skull and head-plate. A bone wax and mineral oil mixture was used to cover the craniotomy. A plastic cap was used to protect the microdrive/silicon probe assembly.

### Anatomy

On the last day of recording, the animals were anesthetized at the end of the recording and perfused transcardially with 4% paraformaldehyde in phosphate buffer. The brain was removed and kept overnight in 4% paraformaldehyde solution. Coronal sections (100 µm thick) were sliced using a vibratome and were mounted on slides using Vectashield mounting medium with DAPI. Images of DAPI and DiI fluorescence were acquired separately with a Nikon FN1 microscope equipped for fluorescence imaging. The DG and the electrode signals were isolated and visualized in 3D using custom MATLAB routines.

### Behavioral control and data acquisition

The forward and backward movement increments of the treadmill were monitored using two pairs of LEDs and photosensors that read patterns on a disc coupled to the treadmill wheel, while the zero position was implemented by an LED and photosensor coupling that detected a small hole on the belt. From these signals, the mouse position was implemented in real time by an Arduino board (Arduino Uno, arduino.cc), which also controlled the valves for the reward delivery. Position, time and reward information from the Arduino board was sent via USB serial communication to a computer and recorded with custom LabView (National Instruments) programs.

Neurophysiological signals were continuously acquired at 30 kHz on a 250-channel recording system (Intan Technologies, RHD2132 amplifier board with RHD2000 USB Interface Board and a custom LabView interface). The wideband signals were digitally high-pass filtered (0.8-5 kHz) offline for spike detection and low-pass filtered (0-500 Hz) and downsampled to 1000 Hz to extract local field potentials. Spikes from each session and each shank of the silicon probe were clustered separately with automatic algorithms ^(^Henze and Buzsáki, 2007) followed by manual adjustments in custom MATLAB routines implementing spike autocorrelation, cross-correlation and cluster isolation statistics. Only clusters with well-defined cluster boundaries and clear refractory periods were included in the analyses (Kadir et al., 2014).

### Estimation of cell position relative to the shank

To estimate the position of a cell relative to the recording sites on a shank, we assumed that the amplitude of the spike signals are attenuated as 1/d^2^ (see note below), where d is the distance of the site to the cell soma, such that the amplitude measured at a given site is:

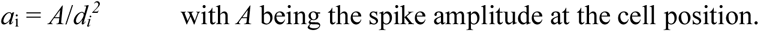

For the many recording sites on one shank, this means that:

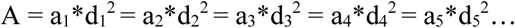

Therefore, to estimate the position of a cell, we simply searched for the position at which these conditions were fulfilled. To do this, the volume around each shank was divided into 1µm^3^ pixels, and for each pixel, we computed the Euclidean distances to each recording site. Then, we defined a value S such that:

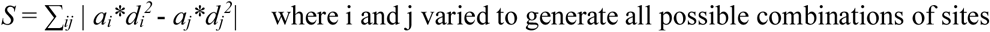

The pixel with the smallest value of *S* was defined as the cell position.

Note: Electric potential of dipoles are attenuated as 1/*d*^*2*^ but as 1/*d* for monopoles. We tested this method using either form and found the resulting cell positions to be very similar.

### Spike gamma phase

The LFP from a channel in the hilus was bandpass filtered between 30-80 Hz. A vector of instantaneous phase was derived using the Hilbert transform. The gamma phase of each spike was interpolated from the vector of the instantaneous phase.

### Gamma coupling index

The ‘gamma coupling index’ captures the coupling of LFP gamma power to cell activity during animal immobility. For each cell, the average gamma power of a hilar LFP was computed for windows within (−10 to +10 ms) and outside (+40 to +100 ms) epochs of maximal firing activity, and the gamma coupling index was defined as the difference between the two windows divided by the sum of the two windows. The 100 largest peaks of the smoothed (using a 20 ms half-width Gaussian kernel) instantaneous firing rate vector were used as epochs of maximal firing.

### Implementation of single neuron firing rate vector

The length of the belt was divided into 100 pixels. To generate a vector of firing rates, the number of spikes discharged in each pixel was divided by the time the animal spent in the pixel. The firing rate vector was smoothed by convolving a Gaussian function (15 cm half-height width).

### Place field emergence and extinction

For a given place field, the mean firing rate in a 10-cm window enclosing the field was calculated for each trial, producing a vector of firing rates. The vector was smoothed using a 3-trial half-width Gaussian kernel. The emergence of a place field corresponded to the time point when the firing rates increased and exceeded 20% of the vector peak value, while the extinction of a field corresponded to a decrease below 20% of the vector peak value.

### Statistical analysis

All statistical analyses were performed in MATLAB (MathWorks). The number of animals and the number of recorded cells were similar to those generally employed in the field. The analysis of variance (ANOVA) was used to test mean differences between groups larger than two. Student t-tests were used to test the sample mean. Correlations were computed using Pearson’s correlation coefficient.

## Data availability

The data that were collected for this study are available upon reasonable request.

## Author contribution

S.K. and S.R. designed the project and performed experiments and modeling. S.K. analyzed the data. D.J. contributed to experiments and spike sorting. S.K. and S.R. wrote the manuscript.

## Acknowledgements

This work was supported by the Korea Institute of Science and Technology Institutional Program (Project No. 2E27850) and Institute for Basic Science, grant IBS-R015-D1.

## Additional information

Competing Interests: The authors declare that they have no competing interests.

## References

Aimone, J.B., Wiles, J., and Gage, F. H. (2006). Potential role for adult neurogenesis in the encoding of time in new memories. Nat. Neurosci. 9, 723–727.

Amaral, D.G., Ishizuka, N., and Claiborne, B. (1990). Neurons, numbers and the hippocampal network. Prog. Brain Res. 83, 1–11.

Amaral, D.G., and Lavenex, P. (2007). Hippocampal neuroanatomy. Andersen, P., Morris, R., Amaral, D., Bliss, T., and O’Keefe, J., ed. The Hippocampus book. (New York: Oxford University Press), pp. 37–114.

Barthó, P., Hirase, H., Monconduit, L., Zugaro, M., Harris, K.D., and Buzsáki, G. (2004). Characterization of neocortical principal cells and interneurons by network interactions and extracellular features. J. Neurophysiol. 92, 600–608.

Bliss, T.V.P., and Collingridge, G.L. (1993). A synaptic model of memory: long-term potentiation in the hippocampus. Nature 361, 31–39.

Boss, B.D., Peterson, G.M., and Cowan, W.M. (1985). On the number of neurons in the dentate gyrus of the rat. Brain Res. 338, 144–150.

Bragin, A., Jandó, G., Nádasdy, Z., van Landeghem, M., and Buzsáki, G. (1995). Dentate eeg spikes and associated interneuronal population bursts in the hippocampal hilar region of the rat. J. Neurophysiol. 73, 1691–1705.

Buzsáki, G., and Czéh, G. (1981). Commissural and perforant path interactions in the rat hippocampus. Field potentials and unitary activity. Exp. Brain Res. 43, 429–438.

Carpenter, G.A., and Grossberg, S. (1987). A massively parallel architecture for a self-organizing neural pattern recognition machine. Comp. Vis. Graph. Image Process. 37, 54–115.

Chung, J., Sharif, F., Jung, D., Kim, S., and Royer, S. (2017). Micro-drive and headgear for chronic implant and recovery of optoelectronic probes. Scientific Reports 7, 2773.

Danielson, N.B., Kaifosh, P., Zaremba, J.D., Lovett-Barron, M., Tsai, J., Denny, C.A., Balough, E.M., Godberg, A.R., Drew, L.J, and Hen, R. et al. (2016). Distinct contribution of adult-born hippocampal granule cells to context encoding. Neuron 90, 101–112.

Danielson, N.B., Turi, G.F., Ladow, M., Chavlis, S., Petrantonakis, P.C., Poirazi, P., and Losonczy, A. (2017). In vivo imaging of dentate gyrus mossy cells in behaving mice. Neuron 93, 552–559.

Deshmukh, S.S., and Knierim, J.J. (2011). Representation of non-spatial and spatial information in the lateral entorhinal cortex. Front. Behav. Neurosci. 5, 69.

Diamantaki, M., Frey, M., Berens, P., Preston-Ferrer, P., and Burgalossi, A. (2016). Sparse activity of identified dentate granule cells during spatial exploration. eLife 5, 1109.

Fattahi, M., Sharif, F., Geiller, T., and Royer, S. (2018). Differential representation of landmark and self-motion information along the CA1 radial axis: self-motion generated place fields shift toward landmarks during septal inactivation. J. Neurosci. 38, 6766–6778.

Geiller, T., Fattahi, M., Choi, J.S., and Royer, S. (2017). Place cells are more strongly tied to landmarks in deep than in superficial CA1. Nat. Commun. 8, 14531.

Gonçalves J.T., Schafer S.T., and Gage F.H. (2016). Adult neurogenesis in the hippocampus: from stem cells to behavior. Cell 167, 897–914.

GoodSmith, D., Chan, X., Wang, C., Kim, S.H., Song, H., Burgalossi, A., Christian, K.M., and Knierim, J.J. (2017). Spatial representations of granule cells and mossy cells of the dentate gyrus. Neuron 93, 677–690.

Hafting, T., Fyhn, M., Molden, S., Moser, M.B., and Moser, E.I. (2005). Microstructure of a spatial map in the entorhinal cortex. Nature 436, 801–806.

Hainmueller, T., and Bartos, M. (2018). Parallel emergence of stable and dynamic memory engrams in the hippocampus. Nature 558, 292–296.

Harris, K. D., Henze, D. A., Csicsvari, J., Hirase, H., and Buzsáki, G. (2000). Accuracy of tetrode spike separation as determined by simultaneous intracellular and extracellular measurements. J. Neurophysiol. 84, 401–414.

Henze, D.A., and Buzsáki, G. (2007). Hilar mossy cells: functional identification and activity in vivo. Prog. Brain Res. 163, 199–216.

Henze, D.A., Wittner, L., and Buzsaki, G. (2002). Single granule cells reliably discharge targets in the hippocampal CA3 network in vivo. Nat. Neurosci. 5, 790–795.

Hoydal, O.A., Skytoen, E.R., Andersson, S.O., Moser, M.B., and Moser, E.I. (2019). Object-vector coding in the medial entorhinal cortex. Nature 568, 400–404.

Hunsaker, M.R., Rosenberg, J.S., and Kesner, R.P. (2008). The role of the dentate gyrus, CA3a,b, and CA3c for detecting spatial and environmental novelty. Hippocampus 18, 1064–1073.

Jung, D., Kim, S., Sariev, A., Sharif, F., Kim, D., and Royer, S. (2019). Dentate granule and mossy cells exhibit distinct spatiotemporal responses to local change in a one-dimensional landscape of visual-tactile cues. Sci. Rep. 9, 9545.

Kadir, S.N., Goodman, D.F.M., and Harris, K.D. (2014). High-dimensional cluster analysis with the masked EM algorithm. Neural Comput. 26, 2379–2394.

Kesner, R.P., Taylor, J.O., Hoge, J., and Andy, F. (2015). Role of the dentate gyrus in mediating object-spatial configuration recognition. Neurobiol. Learn. Mem. 118, 42–48.

Knierim, J.J., and Neunuebel, J.P. (2016). Tracking the flow of hippocampal computation: Pattern separation, pattern completion, and attractor dynamics. Neurobiol. Learn. Mem. 129, 38–49.

Larimer, P., and Strowbridge B.W. (2010). Representing information in cell assemblies: persistent activity mediated by semilunar granule cells. Nat. Neurosci. 13, 213–222.

Lee, J.W., Kim, W. R., Sun, W., and Jung, M.W. (2012). Disruption of dentate gyrus blocks effect of visual input on spatial firing of CA1 neurons. J. Neurosci. 32, 12999–13003.

Leutgeb, J.K., Leutgeb, S., Moser, M.B., and Moser, E.I. (2007). Pattern separation in the dentate gyrus and CA3 of the hippocampus. Science 315, 961–966.

Liu, X., Ramirez, S., Pang, P.T., Puryear, C.B., Govindarajan, A., Deisseroth, K., and Tonegawa, S. (2012). Optogenetic stimulation of a hippocampal engram activates fear memory recall. Nature 484, 381–385.

Marín-Burgin, A., Mongiat L.A., Pardi, M.B., and Schinder, A.F. (2012). Unique processing during a period of high excitation/inhibition balance in adult-born neurons. Science 335, 1238–1242.

Marr, D. (1971). Simple Memory: A theory for archicortex. Philos. Trans. R. Soc. Lond. B Biol. Sci. 262, 23–81.

McHugh, T.J., Jones, M.W., Quinn, J.J., Balthasar, N., Coppari, R., Elmquist, J.K., Lowell, B.B., Fanselow, M.S., Wilson, M.A., and Tonegawa, S. (2007). Dentate gyrus NMDA receptors mediate rapid pattern separation in the hippocampal network. Science 317, 94–99.

McNaughton, B.L., Battaglia, F.P., Jensen, O., Moser, E.I., and Moser, M.B. (2006). Path integration and the neural basis of the ‘cognitive map’. Nat. Rev. Neurosci. 7, 663–678.

Norimoto, H., Makino, K., Gao, M., Shikano, Y., Okamoto, K., Ishikawa, T., Sasaki T., Hioki, H., Fujisawa, S., and Ikegaya, Y. (2018). Hippocampal ripples down-regulate synapses. Science 359, 1524–1527.

O’Keefe, J., and Dostrovsky, J. (1971). The hippocampus as a spatial map. Preliminary evidence from unit activity in the freely-moving rat. Brain. Res. 34, 171–175.

Rolls, E.T., and Kesner, R.P. (2006). A computational theory of hippocampal function, and empirical tests of the theory. Prog. Neurobiol. 79, 1–48.

Rolls, E.T., Stringer, S.M., and Elliot, T. (2006). Entorhinal cortex grid cells can map to hippocampal place cells by competitive learning. Net. Comput. Neural Syst. 17, 447–465.

Royer, S., and Paré, D. (2003). Conservation of total synaptic weight through balanced synaptic depression and potentiation. Nature 422, 518–522.

Royer, S., Sirota, A., Patel, J., and Buzsáki, G. (2010). Distinct representations and theta dynamics in dorsal and ventral hippocampus. J. Neurosci. 30, 1777–1787.

Royer, S., Zemelman, B.V., Losonczy, A., Kim, J., Chance, F., Magee, J.C., and Buzsáki, G. (2012). Control of timing, rate and bursts of hippocampal place cells by dendritic and somatic inhibition. Nat. Neurosci. 15, 769–775.

Scharfman, H.E. (1995). Electrophysiological evidence that dentate hilar mossy cells are excitatory and innervate both granule cells and interneurons. J. Neurophysiol. 74, 179–194.

Scharfman, H.E. (2016). The enigmatic mossy cell of the dentate gyrus. Nat. Rev. Neurosci. 17, 562–575.

Schmidt-Hieber, C., Jonas, P., and Bischofberger, J. (2004). Enhanced synaptic plasticity in newly generated granule cells of the adult hippocampus. Nature 429, 184–187.

Senzai, Y., and Buzsáki, G. (2017). Physiological properties and behavioral correlates of hippocampal granule cells and mossy cells. Neuron 93, 691–704.

Si, B., and Treves, A. (2009). The role of competitive learning in the generation of DG fields from EC inputs. Cogn. Neurodyn. 3, 177–187.

Treves, A., and Rolls, E.T. (1994). Computational analysis of the role of the hippocampus in memory. Hippocampus 4, 374–391.

Turrigiano, G.G., Leslie, K.R., Desai, N.S., Rutherford, L.C., and Nelson, S.B. (1998). Activity-dependent scaling of quantal amplitude in neocortical neurons. Nature 391, 892–896.

van Dijk, R.M. Huang, S.H., Slominaka, L., and Amrein, I. (2016). Taxonomic separation of hippocampal networks: principal cell populations and adult neurogenesis. Front. Neuroanat. 10, 22.

Vivar, C., Potter, M.C., Choi, J., Lee, J.Y., Stringer, T.P., Callaway, E.M., Gage, F.H., Suh, H., and van Praag, H. (2012). Monosynaptic inputs to new neurons in the dentate gyrus. Nat. Commun. 3, 1107.

Vyazovskiy, V.V., Cirelli, C., Pfister-Genskow, M., Faraguna, U., and Tononi, G. (2008). Molecular and electrophysiological evidence for net synaptic potentiation in wake and depression in sleep. Nat. Neurosci. 11, 200–208.

