## Supplementary Figures for "Place cell map genesis via competitive learning and conjunctive coding in the dentate gyrus"

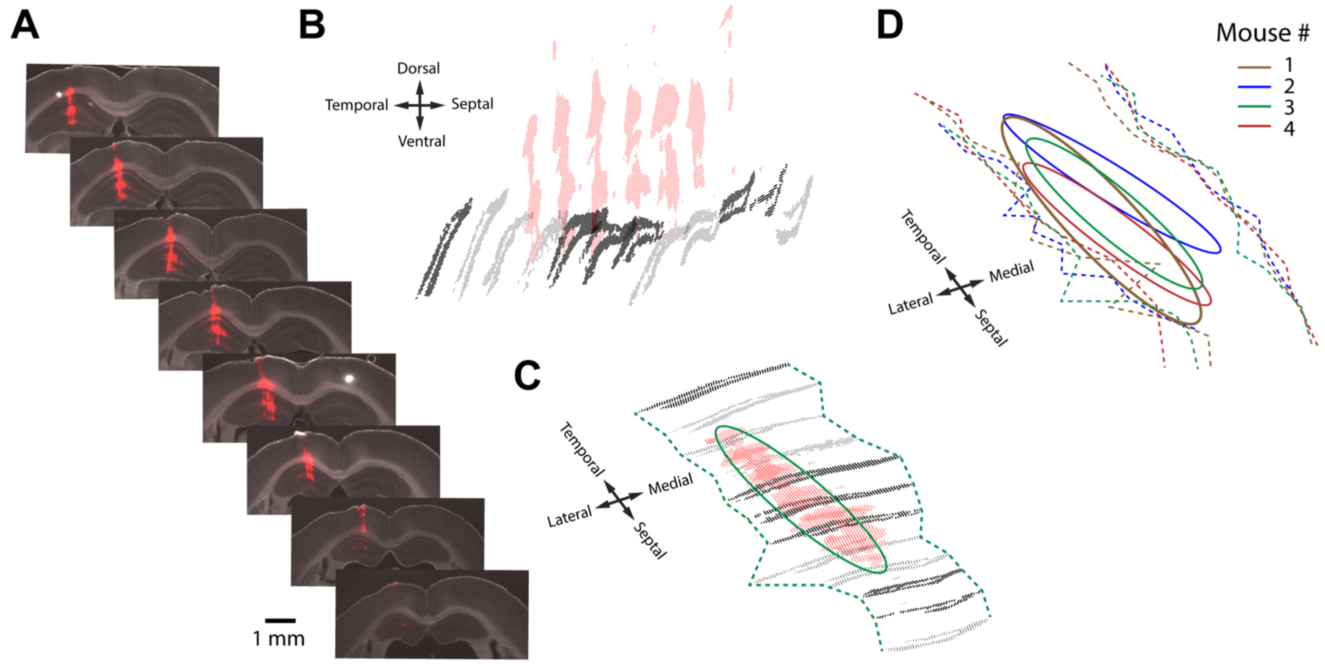


**Figure S1. Histology and electrode positions; related to Figure 1**

A. Example of coronal sections for one animal showing the tracks of the electrodes (red, DiI). Images reconstructed by overlaying DAPI and DiI fluorescence images.

B. Granule cell layers (black dots) and electrode tracks (red dots) were detected for each coronal section using custom MATLAB routines, and 3D reconstructions were implemented for each mouse.

C. The 3D images were rotated perpendicular to the electrode tracks to estimate the electrode positions (ellipsoid) relative to the lateral/medial edges of the granule cell layer (dashed lines).

D. Summary diagram of electrode positions for all mice.

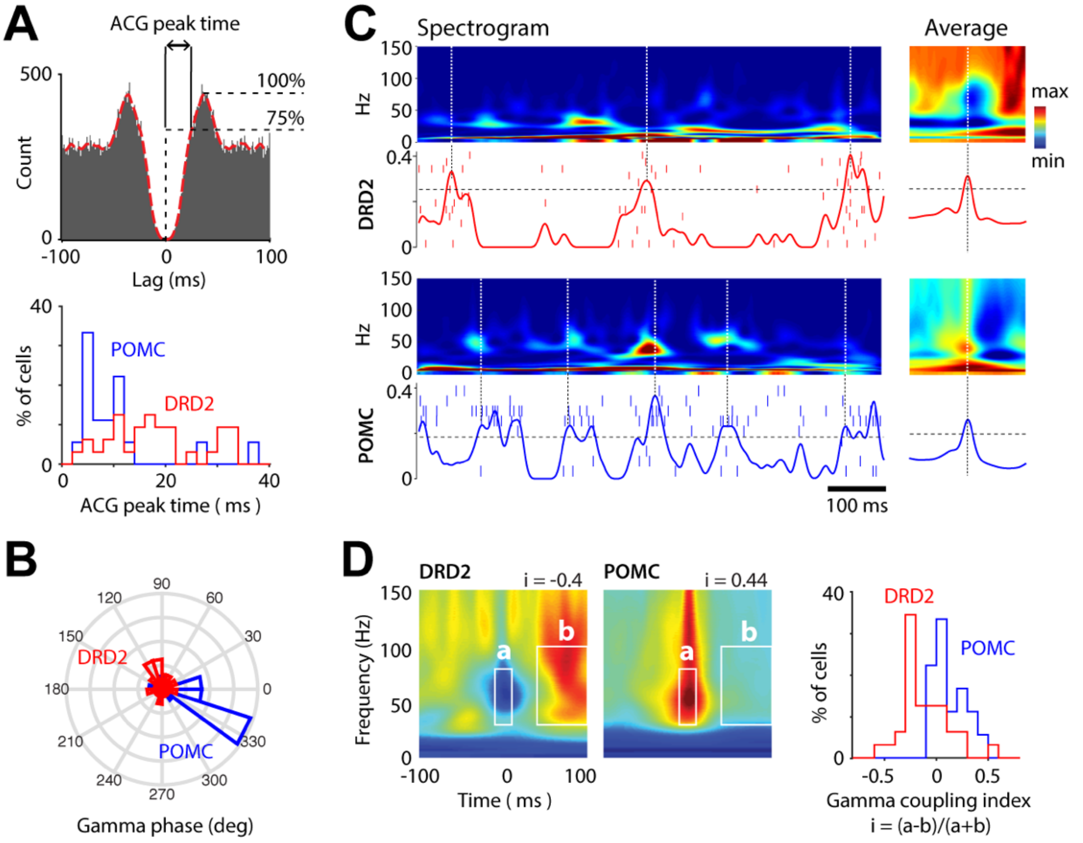


**Figure S2. Spike features of light-excited cells in DRD2 and POMC mice; related to Figure 1**

A. *Upper*, ACG of a cell example and illustration of the ACG refractory gap measure (defined as the duration for spike ACGs to reach 75% of peak value). Lower, distribution of ACG refractory gap for light-excited cells in DRD2 (red) and POMC (blue) mice.

B. Distribution of the mean gamma phase for DRD2 (red) and POMC (blue) light-excited cells.

C. Coupling of cell activity with gamma power. Spectrograms (color-coded), spike rasters of light-excited cells (ticks) and average instant firing rate of light-excited cells (lines), for one DRD2 (upper) and one POMC mouse (lower). Events of strong population activity are detected on the instant firing rates (using a threshold equal to 10% of the largest event, dash lines, left) to generate event-triggered spectrogram averages (right). Note that the activity of light-excited cells is inversely correlated with gamma power for DRD2 and POMC.

D. Gamma coupling indexes of individual cells. *Left*, spike-triggered spectrogram averages for one DRD2 and one POMC light-excited cell. To compute the gamma coupling index i, the mean power in regions a (-10 to 10 ms, 30 to 80 Hz) and b (30 to 100 ms, 30 to 100 Hz) were calculated, and then i = (a-b)/(a+b). *Right*, distribution of gamma coupling indexes for DRD2 and POMC light-excited cells.

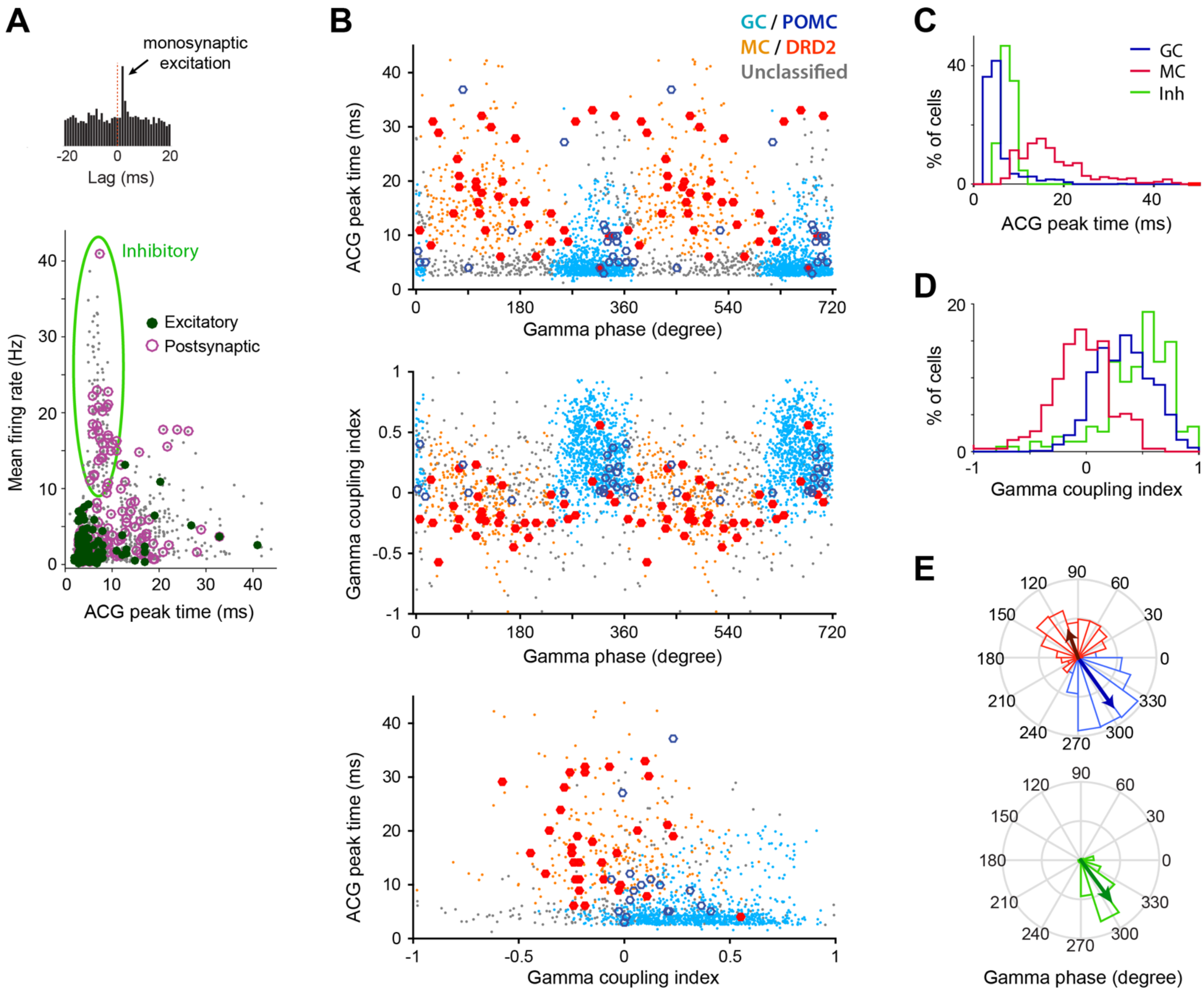


**Figure S3. Identification of putative GCs and MCs; related to Figure 1**

A. Mean firing rate (during animal immobility) versus ACG refractory gap for all cells (*gray*) and a subset of cells identified as presynaptic excitatory (*green*) and postsynaptic (*purple*) cells from a large peak at monosynaptic latency <3 ms in the cross-correlogram of a neuron pair (*inset*). *Green ellipsoid*, putative inhibitory interneurons segregated by high firing rate, short ACG refractory gap and lack of identified excitatory neurons.

B. Scatter plots for spike gamma phase, ACG refractory gap and gamma coupling index of all cells, except the putative inhibitory cells identified in (A). Putative MCs (*orange dots*) and GCs (*light blue dots*) are identified by overlap with DRD2 (*red filled circle*) and POMC (*blue circle*) light-excited cells.

C-E. Distribution of ACG refractory gap (C), gamma coupling index (D) and spike gamma phase (E) for putative GCs (*blue*), MCs (*red*) and inhibitory cells (*green*).

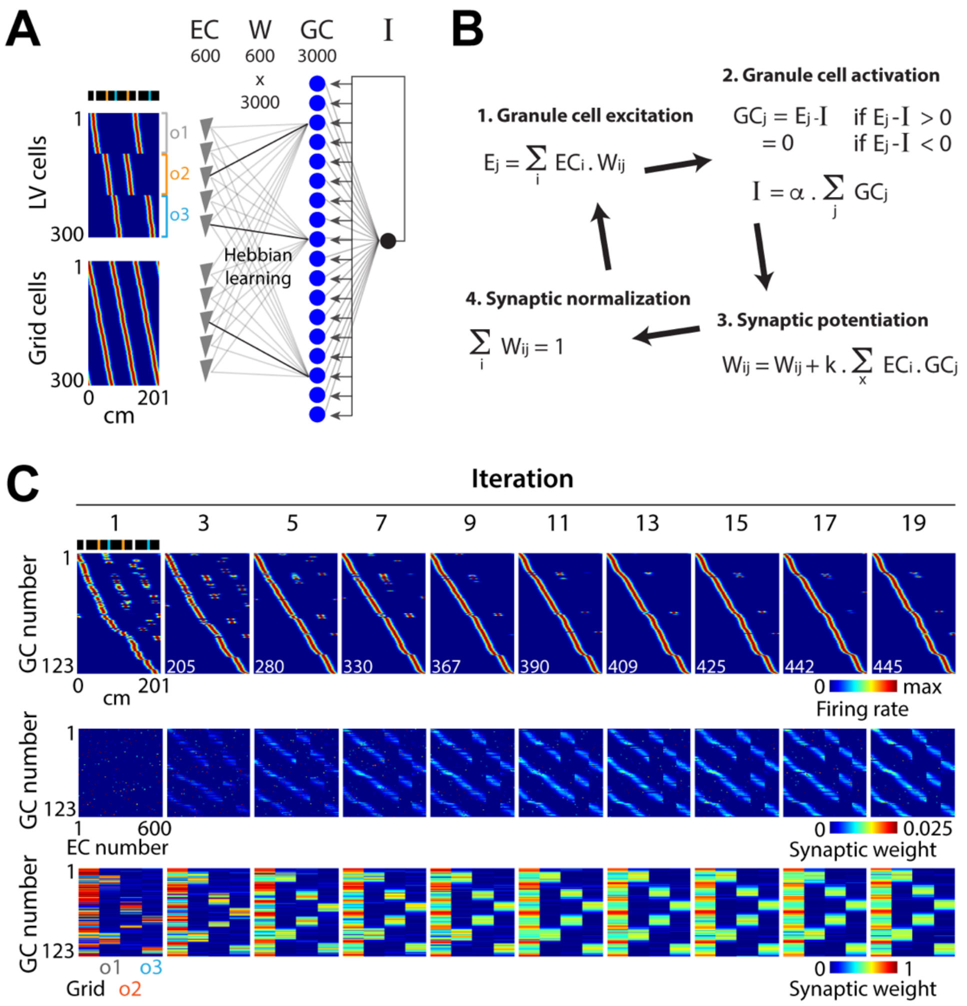


**Figure S4. Competitive learning reproduces the increase in GC single field representations; related to Figure 7**

A. Model architecture. 3000 GCs receive excitatory inputs from 300 LV cells and 300 grid cells from the EC, and are subjected to feedback inhibition. The EC-to-GC synaptic weight matrix is referred to as Wij. The threshold used as feedback inhibition is referred to as I. *Color coded*, rate maps of EC LV cells (upper) and grid cells (lower). LV cells encode various distances to landmarks, while grid cells have various spatial phases and the same periodicity as periodic cells.

B. All operations were executed during one model iteration. 1) The excitation Ej received by the GCj in a given position. 2) The levels of GCj activation and feedback inhibition I in that position. The value of I is estimated numerically by finding the value among a range of I values that best satisfies the two relations. 3) The potentiation of synaptic weights, proportional to the level of EC-GC co-firing throughout the belt. 4) The normalization of synaptic weights.

C. Transformation of spatial representations and synaptic weights across iterations. *Upper*, color-coded representation of rate maps for active GCs across iterations. Active cells have a mean activity > 0 and are sorted according to firing field positions. Note the increase in the number of active GCs (lower left numbers) and the transformation of representations from multiple to single place fields. *Middle*, color-coded representation of synaptic weights for active GCs across iterations, using the same GC ordering as above. Each pixel of the matrices corresponds to one synapse. Note the initial sparse connectivity and the progressive rise in synaptic weights. *Lower*, for each active GC, the color-coded representation of the total synaptic weight contributed by grid cells and by each population of LV cells (encoding a specific landmark) is shown across iterations, using the same GC ordering as above. Note that synaptic inputs initially originate from one source and are progressively redistributed equally among grid cells and LV cells.

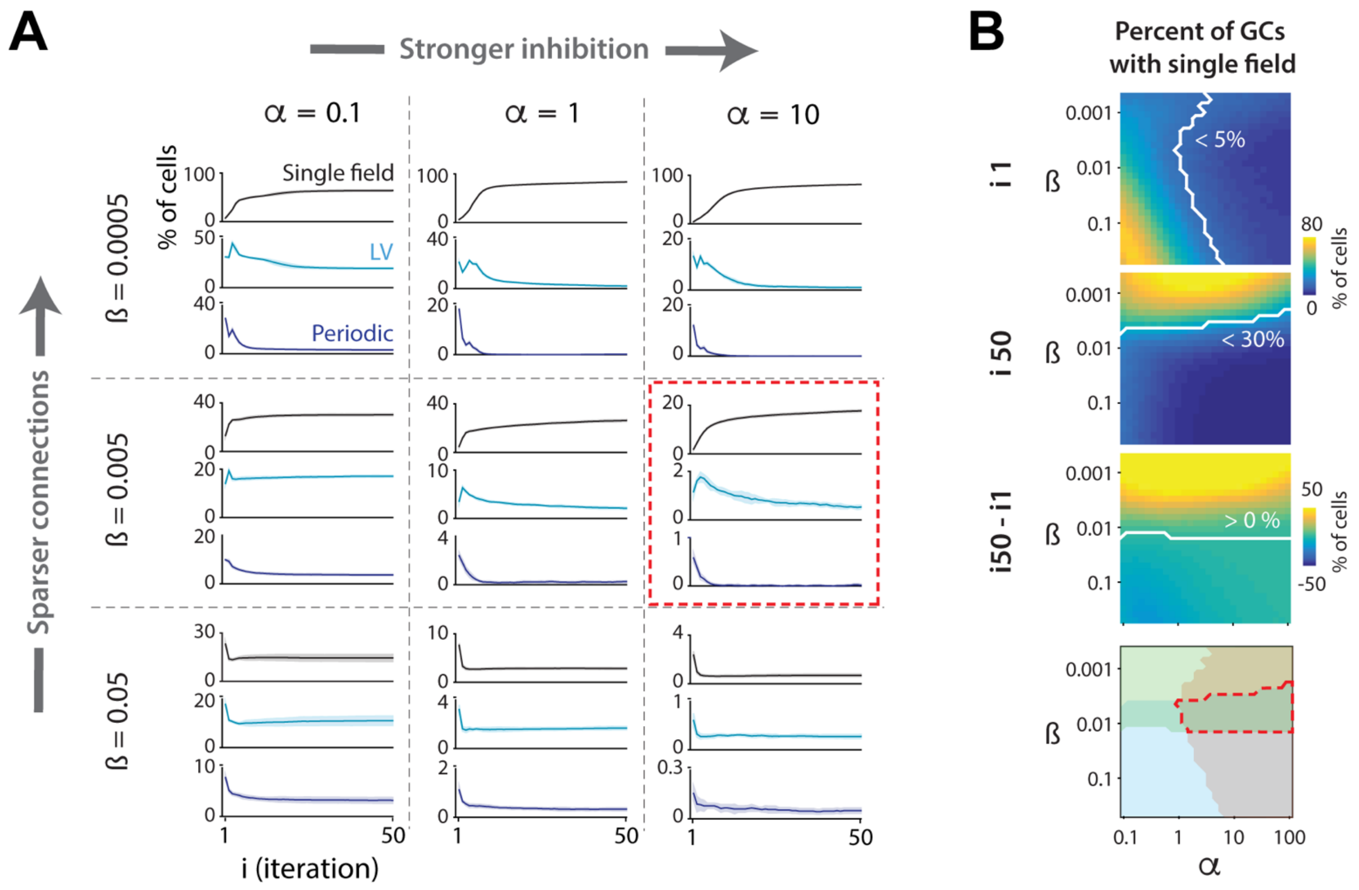


**Figure S5. Impact of inhibition and sparsity of initial inputs; related to Figure 7**

A. Proportion of single (black), LV (light blue) and periodic (dark blue) firing field representations, across iterations, for different values of the parameters alpha and beta, used respectively to adjust the strength of inhibition and the initial sparsity of connections. The red rectangle indicates the conditions that best reproduce the experimental findings.

B. *Top*, color-coded representation of the proportion of GCs exhibiting a single place field in the first model iteration (i1), for the 28 alpha and 28 beta values tested. The pixels on the right of the white line have proportions <5% and are considered compatible with the experimental data. *Second from top*, the same as above for the 50th model iteration (i50). The pixels below the white line have proportions <30% and are considered compatible with the experimental data. *Third from top*, the difference between the 2 matrices above (i50-i1). The pixels above the white line have values >0 (i.e., single field are increased) and are considered compatible with the experimental data. *Bottom*, overlay of the compatible zones defined above. The region of overlap between the zones (red dashed line) provides the possible alpha/beta values to reproduce experimental trends.

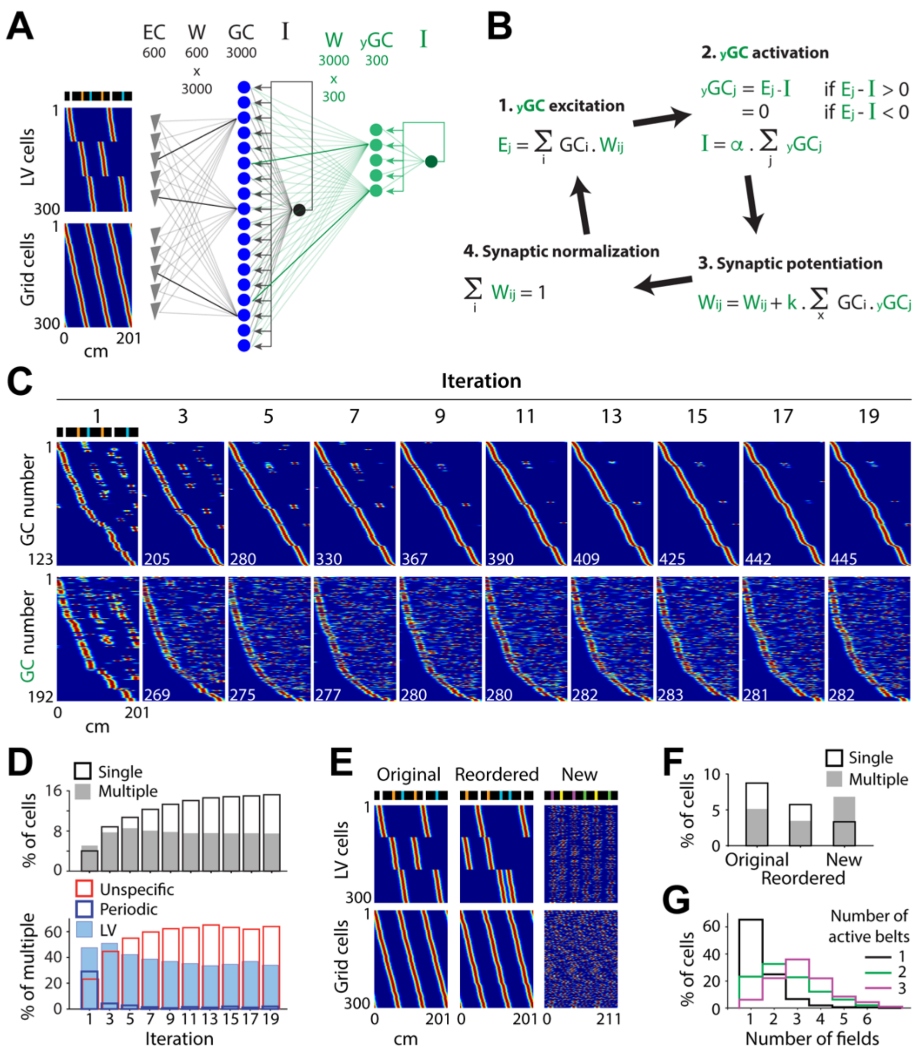


**Figure S6. Modeling of GC multiple unspecific firing fields; related to Figure 7**

A. Model architecture. Three hundred ‘immature’ GCs (green) that receive inputs from the 3000 ‘mature’ GCs and that are subjected to a low level of feedback inhibition are added to the previous model.

B. The same operations as those for mature GCs are executed for immature GCs in each iteration.

C. Color-coded representation of rate maps for active mature GCs (upper) and active immature GCs (lower), across iterations. Active cells have a mean activity > 0 and are sorted according to firing field positions. Note that immature GC representations initially resemble mature GC representations and then become multiple unspecific fields.

D. *Upper*, proportion of GCs (irrespective of type) with a single field (black) and multiple fields (grey), across iterations. *Lower*, proportion of LV, periodic and unspecific GCs, among multiple field GCs, across iterations. Note the similarity with experimental trends (Figure 1C).

E. The rate map of EC LV cells (upper) and grid cells (lower) used to model the original (left), reordered (middle) and new (right) belts. To simulate the reordered belt, the firing fields of EC LV cells are moved to the new location of the landmarks. To simulate the new belt, EC LV and grid cells are randomly assigned to new landmarks and spatial grid phases, respectively.

F. Fraction of GCs with a single (black) and multiple (gray) place fields in each belt. Note the similarity with experimental trends (Figure 3B).

G. The distribution of the number of fields per cell (using the belt with the largest number of fields) for the groups of GCs that are active in 1 (black), 2 (green) and 3 (purple) belts. Note the similarity with experimental trends (Figure 3D).

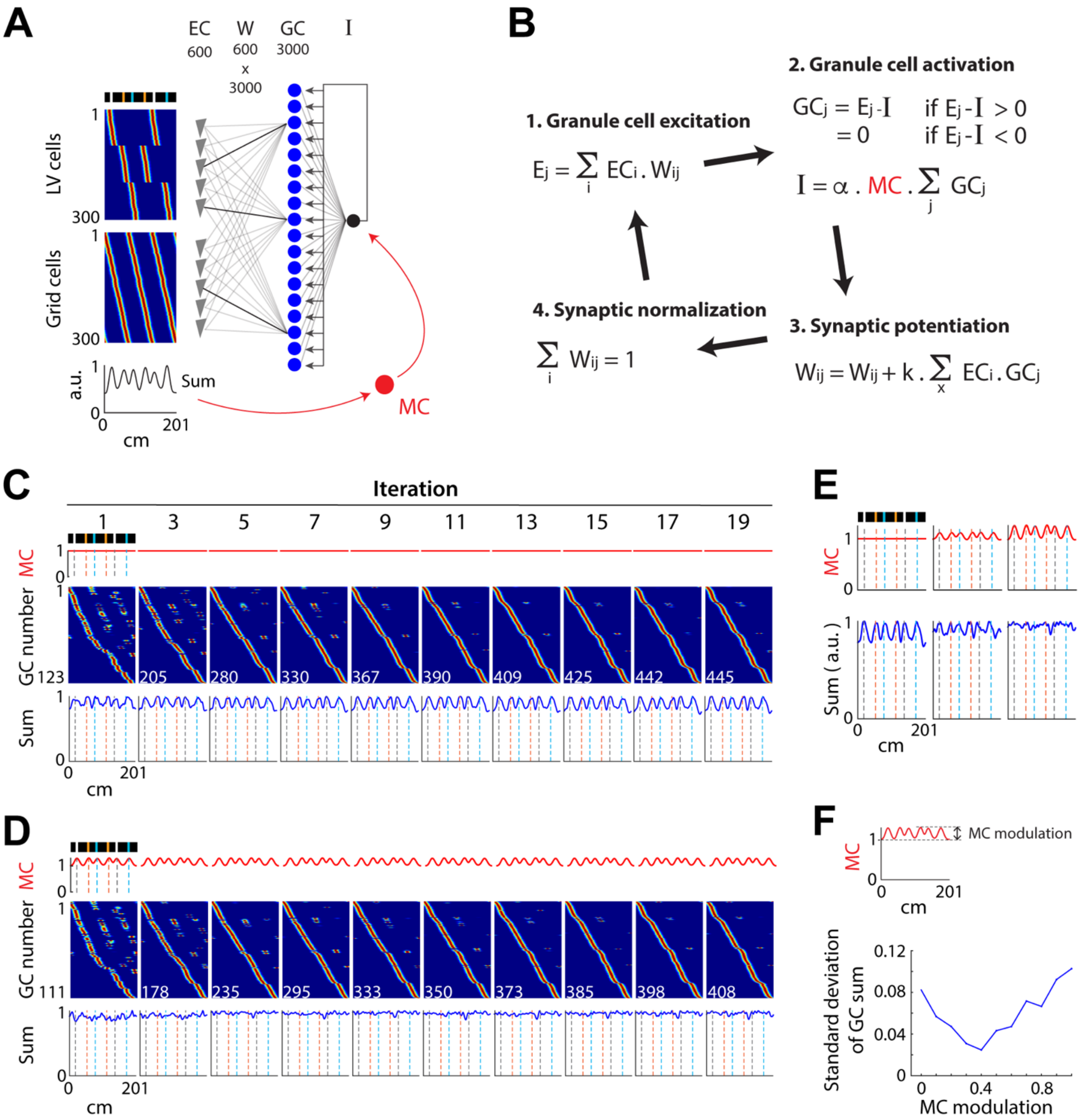


**Figure S7. Modeling the contribution of MC feedforward inhibition; related to Figure 7**

A. Model architecture. MC feedforward inhibition is added to the previous model. The spatial modulation of MC activity is assumed to be proportional to the dynamic range of EC average activity.

B. All operations were executed during one model iteration. The only difference between this model and the previous model is that the strength of the inhibition is modulated throughout space by the factor MC.

C. In the absence of MC modulation, the average activity of GCs is increased in landmark positions. MC activity (top), GC rate maps (color-coded) and the mean of GC rate maps (bottom), across iterations.

D. With MC modulation, the average activity of GCs is uniform throughout the belt. The same format as c.

E. Effect of the distinct magnitude of MC modulation on the GC mean activity.

F. Spatial fluctuation in the GC mean activity as a function of MC modulation.
